# On the variability of dynamic functional connectivity assessment methods

**DOI:** 10.1101/2023.07.13.548883

**Authors:** Mohammad Torabi, Georgios D. Mitsis, Jean-Baptiste Poline

## Abstract

Dynamic functional connectivity (dFC) has become an important measure for understanding brain function and as a potential biomarker. However, various methodologies have been developed for assessing dFC, and it is unclear how the choice of method affects the results. In this work, we aimed to study the results variability of commonly-used dFC methods. We implemented seven dFC assessment methods in Python and used them to analyze fMRI data of 395 subjects from the Human Connectome Project. We measured the pairwise similarity of dFC results using several similarity metrics in terms of overall, temporal, spatial, and inter-subject similarity. Our results showed a range of weak to strong similarity between the results of different methods, indicating considerable overall variability. Surprisingly, the observed variability in dFC estimates was comparable to the expected natural variation over time, emphasizing the impact of methodological choices on the results. Our findings revealed three distinct groups of methods with significant inter-group variability, each exhibiting distinct assumptions and advantages. These findings highlight the need for multi-analysis approaches to capture the full range of dFC variation. They also emphasize the importance of distinguishing neural-driven dFC variations from physiological confounds, and developing validation frameworks under a known ground truth. To facilitate such investigations, we provide an open-source Python toolbox that enables multi-analysis dFC assessment. This study sheds light on the impact of dFC assessment analytical flexibility, emphasizing the need for careful method selection and validation, and promoting the use of multi-analysis approaches to enhance reliability and interpretability of dFC studies.

## Introduction

Functional Connectivity (FC) has emerged as an important measure for understanding brain function, and as a biomarker with significant potential [1–4]. FC is typically assessed using blood-oxygen-level-dependent (BOLD) signals from functional magnetic resonance imaging (fMRI). FC assessment was initially performed under the assumption that it does not change over time (stationarity). However, growing evidence supports that the dynamic variations of FC (dynamic FC – dFC) play a critical role in brain functional organization and may provide a link between neural dynamics and cognition in healthy and diseased conditions [5–7]. The significance of dFC extends beyond understanding interactions between brain regions. Recent studies have shown that patterns of dFC can reveal differences between healthy and diseased individuals and therefore could be used as a clinical biomarker. In this regard, several studies have reported alterations in dFC patterns in a number of neurological or psychiatric conditions (autism spectrum disorders (ASD): [8]; attention-deficit/hyperactivity disorder (ADHD): [9]; depression: [10]; post-traumatic stress disorder (PTSD): [11]; schizophrenia (SZ): [12]; Parkinson’s disease (PD): [13]; Alzheimer’s disease (AD): [14]).

In recent years, a variety of methodologies for assessing dFC have been developed [4, 15]. As the number of available methods continues to grow, there is an increasing need to comprehensively review these methodologies and examine their relative advantages and disadvantages, as well as converge towards recommendations for their application to experimental data [16]. To date, several studies have reviewed different aspects of these methods and have highlighted the importance of understanding their limitations and underlying assumptions [4, 15–21]. However, only a few of these studies have compared different methods in practice [17, 18], and none have provided a comprehensive comparison of the results yielded by commonly-used dFC assessment methods.

The majority of studies that have applied dFC to various applications have not reported a clear justification for adopting a specific methodology for assessing dFC [8, 9, 14, 17, 22–25]. For example, in clinical applications, some studies have employed sliding window and clustering methods (e.g. PTSD: [11]; PD: [24]; SZ: [16, 23, 26]), while others have preferred Hidden Markov Model (HMM) approaches (e.g. SZ: [27]; PTSD: [28, 29]; mild cognitive impairment (MCI): [30]), or Time-Frequency methods (e.g. ASD: [31, 32]; SZ: [33]). It’s worth noting that very few studies have used multiple methodologies (e.g. Chronic headache: [34]; disorders of consciousness (DOC): [35]). Similarly, in cognitive and behavioral applications, some studies have used sliding window and clustering methods (e.g. task prediction: [22]; cognitive and behavioral flexibility: [36]), some have used CoActivation Patterns (CAP) analysis (e.g. naturalistic stimuli: [37]; multiple tasks: [38]), HMM (e.g. Sleep stage: [39]), and very few have used multiple methods (e.g. working memory task: [40]). A complete list of the reviewed studies can be found in Tables 2, 3, and 4 in Supplementary material. Strikingly, out of the 60 dFC studies that we reviewed, only four considered more than one method to assess the robustness of their results. This is somewhat concerning since the variation in the dFC assessment results due to the choice of the assessment methodology, i.e. its analytical flexibility, has been studied on a relatively limited basis until now [17].

Analytical flexibility has recently become a critical issue in neuroimaging, particularly in fMRI data analysis [41–44]. Several studies have revealed significant variations in the results obtained from multiple independent analyses using the same dataset, highlighting the impact that analytical flexibility may have on scientific findings. For example, a study by [41] revealed considerable variation in the results reported by 70 independent teams using a common dataset, emphasizing the need for performing and comparing multiple analyses on the same dataset, as well as identifying the factors that explain the variation in the obtained results.

Similarly, the analytical flexibility of dFC assessment may greatly influence the reliability of the findings about brain functional organization and its alterations in diseased groups. Although several studies have investigated the reproducibility across individuals and scan sites of dFC patterns at rest [26, 45, 46], no study has fully investigated the analytical flexibility of dFC assessments. It is also unclear how methodological variations compare to the expected biological variations. The need for understanding dFC analytical flexibility in the context of biological variability is a first critical step for result interpretation and for drawing conclusions about brain functional organization and its alterations in diseased groups.Resting-state fMRI data lack a reliable biological reference for quantifying the scale of variability of dFC across assessment methods. It is well known that fMRI data are an indirect measure of the neural activity and are also modulated by physiological and motion processes [47, 48]. This further complicates setting a biological reference. However, dFC variation over time or over subjects can serve as fair biological references. Comparing the variability over methods to dFC temporal variability would yield an interpretable biological scale. In other words, if assessing dFC at one time point across methods results in as much variation as the one found across time points using one method, then variability between methods is as large as the scale of the biological phenomenon that they intend to capture. This approach enabled us to directly compare the magnitude of the effect of methodology choice to the underlying biological phenomenon. An alternative to variability in time is inter-subject variability, which can also help scale the magnitude of variations over method.

In this study we assess the analytical flexibility of dFC estimation by investigating how dFC patterns estimated from the same dataset vary across a large selection of methods. Among the available dFC assessment methods, we have selected seven widely used methods: Co-Activation Patterns (CAP) [49], the clustering method [5], continuous Hidden Markov Models (HMM) [50], discrete HMM [28], the sliding-window method [12], the time-frequency (wavelet-based) method [7], and the window-less method [51].

The selected dFC assessment methods differ in terms of their basic underlying assumptions, ranging from the window-less method, which places a minimal set of assumptions to the HMM-based methods which place the most rigid assumptions [4]. Furthermore, some of these methods are state-based methods, meaning that they assume a finite number of group-level FC spatiotemporal patterns or “FC states” recurring over time (CAP, clustering, continuous HMM, discrete HMM, and window-less), while the rest are state-free methods which do not impose any constraints on the estimated FC patterns (sliding window and time-frequency). Moreover, the examined methods differ with regards to whether they assume locality of neighboring time points. Briefly, locality assumption means that the FC at each time point is inferred from the data points that are close in time (sliding window, time-frequency, clustering, continuous HMM, and discrete HMM), or not (CAP and window-less). The methods also differ in whether they consider the temporal ordering of data points (sliding window, time-frequency, clustering, continuous HMM, and discrete HMM) or disregard this information (CAP and window-less). State-based methods may also vary in terms of assuming smooth transitions between different states (continuous HMM, and discrete HMM) or allowing instantaneous reconfiguration (CAP, clustering, and window-less) [4]. Table1 lists these seven methods along with their number of citations (note that these numbers refer to the particular paper that introduced them, hence they may not be entirely reflective of the overall popularity of each method), year of publication, and assumptions. For the detailed pipeline used for each method, as well as their important hyperparameters, please see Figure1.

**Fig. 1.**
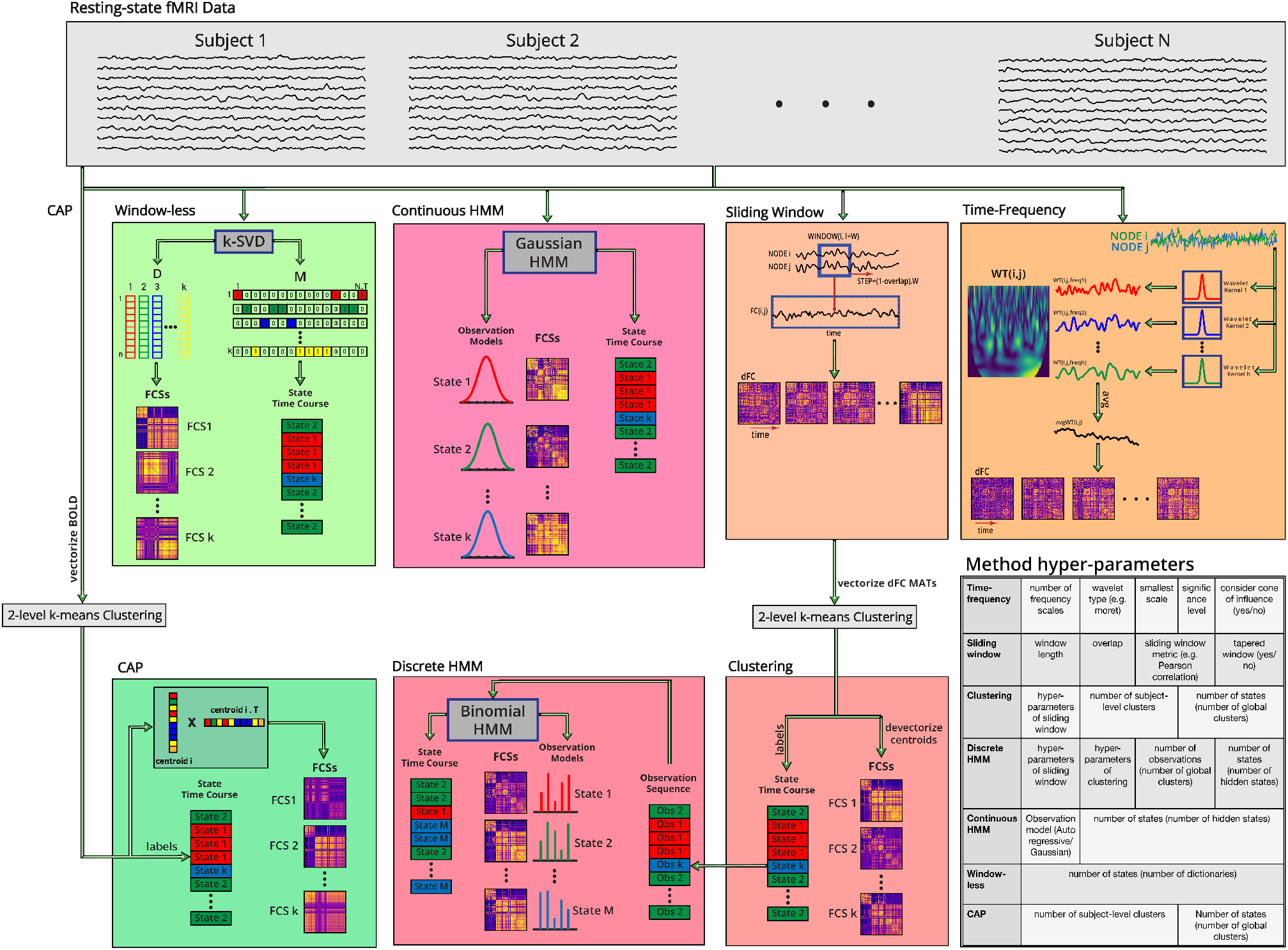
pipeline of the seven dFC assessment methods (time-series of all subjects are concatenated in time dimension for state-based methods prior to finding FC states): **Time-frequency** [7]: Wavelet Transform Coherence (WTC) between each pair of regions is computed after applying the wavelet transform on region time courses. Finally, dFC assessment is performed by averaging WTC over all frequency scales. **Sliding window** [12]: dFC is assessed for each time window using a sliding window which moves over data time points and typically uses Pearson correlation as the metric to quantify FC for each pair of regions. **Clustering** [5]: applies 2-level k-means clustering on the sequence of measured dFC matrices from sliding window method to find the clusters that correspond to different FC states. Cluster centroids are considered as the state FC matrices, and the sequence of clustering labels is considered as the state time course. **Discrete Hidden Markov Model (HMM)** [28]: the state FC matrices and the state time course inferred by clustering are used as the observation sequence of a binomial HMM. The model then finds the hidden states and infers the corresponding state FC matrices and the state time course. **Continuous HMM** [50]: the BOLD data is fed to a Gaussian HMM. The model finds the hidden states and their continuous observation models. It then infers the state FC matrices and the state time course. **Window-less** [51]: a sparse dictionary learning is applied on the BOLD data using a k-SVD model to infer the dictionary matrix (D) and the mixing matrix (M). Next, state FC matrices and the state time course are obtained from D and M respectively. **Co-Activation Pattern (CAP)** [49]: clustering is performed on vectors of BOLD activity at each time point. Next, outer product of each centroid vector and its transpose is calculated and forms the state FC matrix corresponding to that cluster. The sequence of clustering labels of time points is considered as the state time course. Table A shows a list of all implemented dFC assessment methods and their important hyperparameters. Note that the state FC matrices in this figure are denoted using the term ‘FCS.’

**Table 1.**
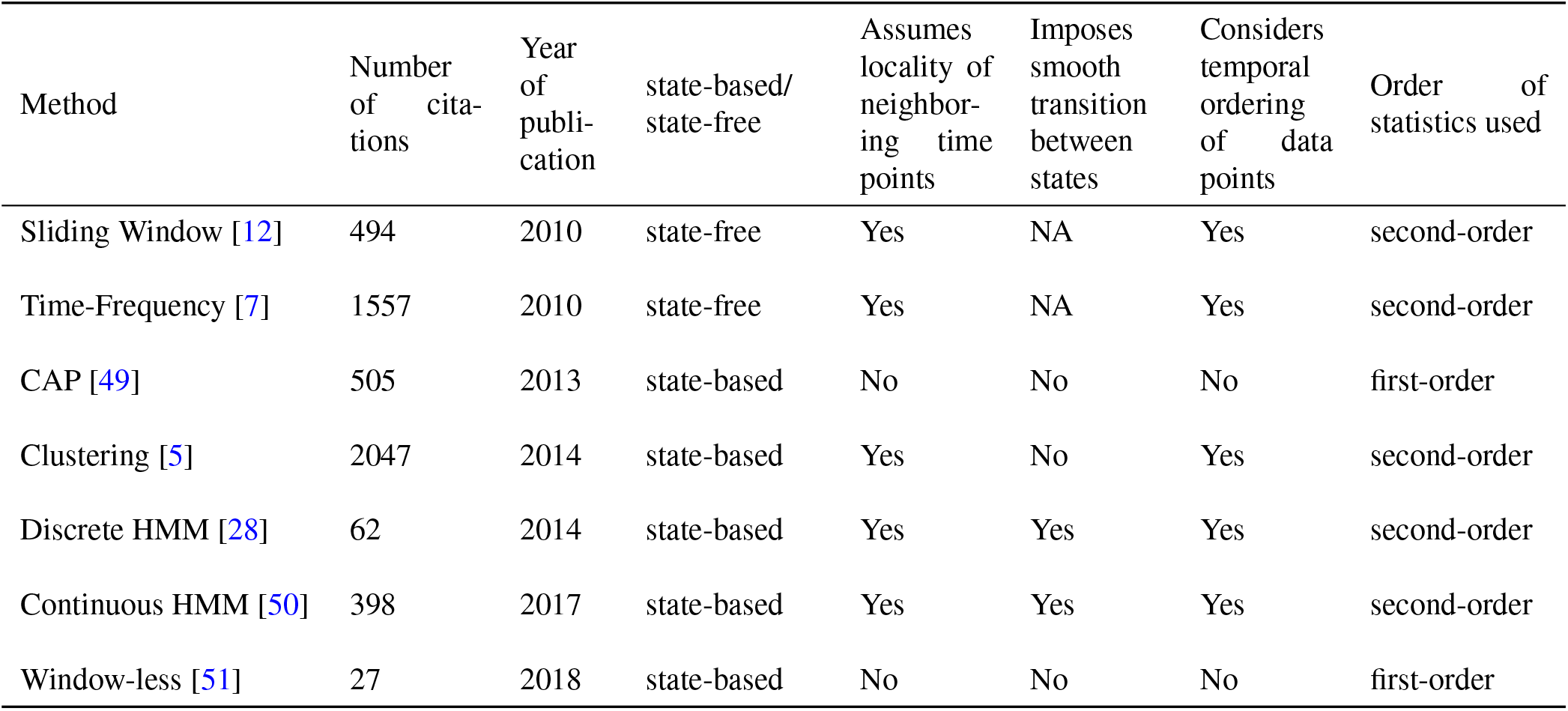
the seven selected dFC assessment methods and their number of citations, year of first publication, and assumptions.

| Method | Number of citations | Year of publication | state-based/<br>state-free | Assumes locality of neighboring time points | Imposes smooth transition between states | Considers temporal ordering of data points | Order of statistics used |
| --- | --- | --- | --- | --- | --- | --- | --- |
| Sliding Window [12] | 494 | 2010 | state-free | Yes | NA | Yes | second-order |
| Time-Frequency [7] | 1557 | 2010 | state-free | Yes | NA | Yes | second-order |
| CAP [49] | 505 | 2013 | state-based | No | No | No | first-order |
| Clustering [5] | 2047 | 2014 | state-based | Yes | No | Yes | second-order |
| Discrete HMM [28] | 62 | 2014 | state-based | Yes | Yes | Yes | second-order |
| Continuous HMM [50] | 398 | 2017 | state-based | Yes | Yes | Yes | second-order |
| Window-less [51] | 27 | 2018 | state-based | No | No | No | first-order |

**Table 2.** Clinical applications of dFC.

| Paper | Application | DFC method applied | Notes |
| --- | --- | --- | --- |
| [31] | ASD | TF | using WTC |
| [32] | ASD | TF | using WTC |
| [8] | ASD | SWC |  |
| [61] | ASD | HMM |  |
| [62] | ASD, AD | SW |  |
| [63] | AD, Dementia risk | SWC |  |
| [14] | AD | SW |  |
| [64] | AD | HMM |  |
| [65] | AD, MCI, SCI | TF | Hilbert transform |
| [30] | MCI | HMM |  |
| [66] | MCI | SW |  |
| [67] | MCI | SW |  |
| [68] | MCI | SW |  |
| [33] | SZ | TF | using WTC |
| [23] | SZ | SWC |  |
| [16] | SZ | SWC |  |
| [69] | SZ | HMM |  |
| [27] | SZ | HMM |  |
| [25] | SZ, BP | SWC |  |
| [13] | PD | SWC |  |
| [24] | PD | SW |  |
| [70] | PD | SWC |  |
| [71] | PD | SWC |  |
| [72] | PD | SW |  |
| [73] | PD | SW |  |
| [74] | PD | CAP |  |
| [28] | PTSD | HMM |  |
| [29] | PTSD | HMM |  |
| [75] | PTSD | SWC |  |
| [11] | PTSD | SW |  |
| [76] | MDD | HMM |  |
| [10] | MDD | SW |  |
| [77] | MDD | SWC |  |
| [9] | ADHD | SWC |  |
| [78] | LBD | HMM |  |
| [79] | Idiopathic generalized epilepsy | SWC |  |
| [80] | Stroke | SWC |  |
| [81] | FTD | SW | with identifying meta-states |
| [35] | DOC | SWC, HMM |  |
| [82] | Aging | SW |  |
| [34] | Chronic headache | SW, TF | using WTC |
| [83] | OCD | SW |  |
| [84] | SZ, ASD, ADHD | SW | 8 datasets ( 2 SZ, 2 ASD, 2 ADHD, 2 fMRI-EGG, 2 fMRI-DTI) |

List of abbreviations: Alzheimer's Disease (AD), Attention-Deficit/Hyperactivity Disorder (ADHD), Autism Spectrum Disorder (ASD), Bipolar (BP), Disorders Of Consciousness (DOC), Diffusion Tensor Imaging (DTI), ElectroEncephaloGram (EEG), FrontoTemporal Dementia (FTD), Lewy Body Dementia (LBD), Mild Cognitive Impairment (MCI), Major Depressive Disorder (MDD), Obsessive-Compulsive Disorder (OCD), Parkinson's Disease (PD), Post-Traumatic Stress Disorder (PTSD), Subjective Cognitive Impairment (SCI), Schizophrenia (SZ), Wavelet Transform Coherence (WTC)

**Table 3.** Cognitive and Behavioral applications of dFC.

| Paper | Application | DFC method applied |
| --- | --- | --- |
| [85] | Dynamic reconfiguration of frontal brain network during executive cognition | SW |
| [86] | Inferring Brain State Dynamics Underlying Naturalistic Stimuli Evoked Emotion Changes | HMM |
| [38] | Across multiple tasks, to investigate anti-correlated functional networks | CAP |
| [37] | Naturalistic stimuli during a TV program | CAP |
| [40] | A sustained 2-back working memory (WM) task compared to rest | CAP, SW |
| [87] | Emotional and cognitive control task | CAP |
| [19] | Distinguish task-induced changes | SWC, MTD, DCC, JC |
| [39] | Sleep stage | HMM |
| [22] | Task prediction | SWC |
| [36] | Cognitive and behavioral flexibility | SWC |

List of abbreviations: Dynamic Conditional Correlation (DCC), Jackknife Correlation (JC), Multiplication of Temporal Derivatives (MTD)

**Table 4.** Other applications of dFC.

| Paper | Application | DFC method applied | Notes |
| --- | --- | --- | --- |
| [88] | Investigating functional organization | SW | developing circuit models |
| [89] | Fusion, investigating functional organization, clinical | SWC | FNC-genomic features, SZ |
| [90] | Fusion | SWC | identifying EEG spectral signatures associated with different FNC states |
| [91] | Investigation | HMM | developmental trajectory across childhood |
| [26] | Clinical, fusion | SWC | SZ, fMRI and structural MRI (grey matter) |
| [92] | Cognitive, clinical | SWC | Sleep deprivation |
| [93] | Investigation, cognitive | SWC | Assess stability of rs-dFC, effect of head motion, sleep state, and imposed task |

List of abbreviations: ElectroEncephaloGram (EEG), Functional Network Connectivity (FNC), Schizophrenia (SZ)

Comparing different methods for assessing dFC poses significant challenges due to several factors. The primary challenge arises from the differences in the underlying assumptions and the mathematical framework of various methods, coupled with the lack of a clear ground truth for measuring the accuracy of the results [4, 19]. Additionally, the distribution of estimated dFC values across methods can differ significantly, precluding the use of common similarity metrics such as the Euclidean distance or Pearson’s correlation. The distribution of dFC values may not necessarily be Gaussian, and these values may considerably vary with regards to their mean and variance (see Figure18 in Supplementary material). Furthermore, the output format of different methods varies, making direct comparisons more challenging. For instance, some methods, such as CAP and window-less, rely on first-order statistics and provide outputs in the form of activity vectors. In contrast, other methods, such as the clustering method, encode dFC as FC matrices, which are second-order statistics. Additionally, state-free methods generate a continuous sequence of FC matrices over time, while state-based methods generate a set of state FC matrices and a state time course.

Comparing dFC methods lacks a general framework. Previous studies that have attempted to compare dFC methods have been limited to specific groups of methods or aspects of dFC. For instance, [17] compared the results obtained by Clustering, CAP, and Phase Synchrony (PS) converted to a discrete set of states, thus limiting the comparisons to state-based methods. To make the results of CAP method comparable to others, they calculated the outer-product of the state mean-activity vectors obtained by this method to construct the corresponding FC matrices. Their comparison criteria were limited to visual comparison of spatial patterns of FC states and a few statistical outcomes derived from FC states, such as fractional occupancy. Similarly, [18] has compared the results obtained by sliding window analysis using ten different correlation metrics. To the best of our knowledge, no studies have comprehensively compared results across the most common dFC assessment methods(state-based and state-free).

To address these challenges, we developed a comprehensive comparison framework applicable to state-based and state-free methods. This framework compares methods based on spatial, temporal, and overall similarity. We implemented the seven selected dFC assessment methods in a coherent Python package, providing the first open-source implementation of its kind. The seven implemented methods were then used to assess dFC using the resting-state fMRI data of 395 subjects from the Human Connectome Project (HCP) dataset [52, 53]. The outputs were standardized into a common array format to allow for comparison of results from methods with different output formats. The resulting dFC arrays were then compared using the comparison framework. This framework employs different similarity metrics, including non-parametric metrics such as Spearman correlation, to deal with the different distributions that result from different dFC methods. This framework was developed to evaluate the variability of the results using two levels of comparison including subject-level and inter-subject level. Beyond assessing temporal and spatial variability on a subject-level, assessing inter-subject similarity enabled us to evaluate the ability of methods to capture a common biological phenomenon by examining the consistency of their inter-subject correlations. This is particularly relevant for studies focused on identifying population biomarkers. For a complete description of the comparison framework see Section **Analytical Flexibility Assessment** in **Methods and Materials**.

Our findings have important implications for the field of dFC and can guide future studies in several ways. Firstly, by comprehensively comparing seven widely used dFC assessment methods, we shed light on their analytical flexibility and the variability of their results, particularly when compared to the inherent variability of the biological phenomenon they aim to capture. This understanding is crucial for researchers and practitioners when choosing an appropriate method for their specific research question or clinical application. Secondly, our study provides a standardized framework for comparing dFC methods, addressing the challenges posed by their diverse assumptions, mathematical foundations, and output formats. This framework can serve as a basis for future comparative studies and facilitate the development and implementation of new or other dFC assessment methods (e.g. [54, 55]). Lastly, our investigation of inter-subject similarity consistency provides insights into the ability of different methods to capture a common biological phenomenon, which is vital for identifying reliable biomarkers of brain function and disorders.

## Methods and Materials

### Data

The data used in this study are Blood-oxygen-level-dependent (BOLD) time series of resting-state fMRI data of 395 young, and healthy subjects (age range: 22-36 years) from the S1200 release of the 3T Human Connectome Project (HCP) dataset [52, 53], preprocessed as described in [48]. Each subject was scanned during four sessions on two different days. Each day included two 15-min scans, one acquired with left-to-right (LR) phase encoding direction and the other one with right-to-left (RL). In this study, except Figure3a, where we show results from all four sessions, all other main results are shown using data from the first session `Rest1_LR`. Each scan consists of 1200 time points with TR=0.72 sec. We used the HCP FIX-denoised data, which includes additional denoising steps such as de-trending, head motion correction, and denoising via spatial ICA compared to the minimally preprocessed data available in the HCP database. The data were then parcellated into 333 regions of interest (ROIs) using the Gordon parcellation [56], an atlas widely used in the literature. Each of the ROIs belongs to a specific Resting State Network (RSN). Among these 333 ROIs, 47 did not belong to any brain network and therefore were excluded, resulting in 286 ROIs. For computational reasons, we have uniformly downsampled these 286 ROIs to 96 ROIs. Each ROI time series was z-standardized before the analysis.

### State-free Methods

These methods are model free and therefore do not need to be implemented at the group level before dFC assessment. They were applied to each subject separately. See Figure1 for an illustration of state-free methods and their dFC matrices.

#### Sliding Window (SW) [12]

A sliding tapered window of length 44s, or 60 time points (following [5]), and 50 percent overlap was used to find the FC of ROI pairs over time. The tapered window was generated by convolving a rectangle of length 44s with a Gaussian curve with *σ* = 3 TRs following [5] and then moved along 1200 time points with a 30 time points overlap. This procedure resulted in 38 windows and hence a 38×*ROI*×*ROI* dFC matrix. The main hyperparameters of this method are window length, overlap ratio, and tapered or rectangular window.

#### Time-Frequency (TF) [7]

The wavelet transform was performed by applying *k* = 101 wavelet kernels on each ROI time course, resulting in a wavelet time series with *k* frequency scales. Next, the Wavelet Transform Coherence (WTC) was computed between the time courses of each ROI pair for each frequency scale and averaged across all frequency scales following [57]. The resulting pairwise WTC time courses formed the corresponding dFC matrices of shape 1200×*ROI*×*ROI*. Note that following [7], only the WTC values outside the cone of influence were considered for the final dFC matrix. The hyperparameters of this method include number of frequency scales, wavelet type, significance level of coherence magnitude, and considering the cone of influence or not.

#### State-based Methods

Since state-based methods assume a finite number of FC states re-occurring over time and all subjects, they initially need to be applied on a group level to identify the group-level FC states. Each state is represented by a state FC matrix of shape *ROI*×*ROI* which corresponds to the FC pattern at the time points that the state occurs. Next, the fitted models are applied to each individual subject, and individual dFC matrices are obtained. Applying the fitted models on subject-level data assigns each time point to one of the group-level FC states, resulting in a state time course. The state FC matrix of each state is then placed at the time points that belong to that state to build the dFC matrix of that subject. There is still no consensus on the number of brain states. In this study, to make the results more comparable, the number of FC states was assumed to be 12 for all state-based methods following [50]. Figures 5 through 9 in Supplementary material show the state FC matrices obtained by each state-based method. See Figure1 for an illustration of state-based methods and their state FC matrices and state time courses.

#### Co-Activation Patterns (CAP) [49]

The Co-Activation Patterns (CAP) method is a point-process analysis with individual time point time resolution and very few assumptions. The original CAP method used activation thresholding within a seed region and subsequent averaging of the BOLD time series at significant time points [49]. The original method does not directly yield dFC matrices. The extended version of CAP analysis [58] applies k-means clustering directly to BOLD time series. In the present study, for computational reasons, the clustering was performed in two stages following a process similar to [28]. First, we clustered the time points of each subject (vectors of *length* = *ROI*) to find subject-level cluster centroids. Next, we clustered the resulting cluster centroids from all subjects to identify 12 group-level cluster centroids, corresponding to 12 FC states. Finally, we obtained the state FC matrices following the implementation proposed by [17], by calculating the *ROI*×*ROI* outer-product matrix of centroids with themselves. The clustering labels of time points were considered as the state time courses. The shape of the final dFC matrix was 1200×*ROI*×*ROI*. The hyperparameters of this method are the number of subject-level clusters, and the number of FC states.

#### Sliding Window + Clustering (SWC) [5]

This method relies on the dFC matrix assessed using the Sliding Window method. Therefore, we initially assessed the dFC matrix using the Sliding Window method as described above, which had a shape of 38×*ROI*×*ROI*. Next, we considered each time point of the dFC matrix as a sample and vectorized it to a feature vector of length *ROI*× (*ROI* −1)/2. Then, using a two-level k-means clustering procedure similar to the one described for the CAP method, we clustered feature vectors into 12 clusters corresponding to 12 FC states. The FC matrix of each state was then obtained by reshaping the centroid vectors to their original shape of *ROI*×*ROI*. The resulting clustering label sequences were considered as the state time courses. The shape of the final dFC matrix was 38×*ROI*×*ROI*. The hyperparameters of this method are those of sliding window method, number of subject-level clusters, and number of FC states.

#### Continuous Hidden Markov Model (CHMM) [50]

In this method, a continuous Hidden Markov Model (continuous HMM or CHMM) with a Gaussian observation model was used. BOLD time series from resting-state fMRI data were directly used as the continuous observation sequences of the HMM. Subsequently, 12 hidden states were identified, each corresponding to a FC state. Each hidden state was represented by a multivariate Gaussian Model. The mean and covariance of each hidden state were inferred by fitting the model on the concatenated BOLD time series of all subjects.

Next, the fitted model was applied on the subject-level data to infer the sequence of hidden states for each subject. The inferred sequence was then used as the state time course and the covariance matrix of each hidden state was considered as the corresponding state FC matrix. The shape of the final dFC matrix was 1200×*ROI*×*ROI*. The only hyperparameters of this method are the observation model type (e.g. Gaussian, or Autoregressive), and number of FC states.

#### Discrete Hidden Markov Model (DHMM) [28]

For this method, a discrete Hidden Markov Model (discrete HMM or DHMM) with a binomial observation model was used. Since this method is based on the results of the clustering method, we initially assessed dFC matrices using the clustering method, which had a shape of 38×*ROI*×*ROI*. The number of FC states assumed for this initial clustering analysis was equal to the number of observations assumed for the discrete HMM method, and not the number of final FC states. Furthermore, the DHMM used here requires a discrete observation sequence. Therefore, the state time courses obtained by the clustering method were used as the discrete observation sequences of the discrete HMM. The HMM was then fitted to the observation sequences to identify 12 hidden states corresponding to the 12 FC states. The resulting hidden state sequences were used as the state time courses and the FC matrix of each FC state was obtained by averaging all FC matrices assigned to that FC state in the dFC matrices obtained by the clustering method. The shape of the final dFC matrix was 38×*ROI*×*ROI*. The hyperparameters of this method include hyperparameters of the clustering method, number of observations, and number of FC states.

#### Window-less (WL) [51]

This method relies on estimating the dominant linear patterns in the sample space of all BOLD time series values. Each dominant linear pattern corresponds to one of the FC states. To estimate the dominant linear patterns, a sparse dictionary learning algorithm, the `k-SVD` algorithm [59], was applied to the group-level data. Dominant linear patterns are each represented by a dictionary element. The 12 dictionary elements corresponding to the 12 FC states were stored in the dictionary matrix *D*. Each time point was approximated by a linear combination of these dictionary elements. The coefficients of the linear combinations for each time point were stored in rows of the mixing matrix *M*. A hard sparsity constrain was applied to the algorithm so that it assigns one and only one dictionary to each time point, suggesting that rows of *M* were sparse. The state FC matrices were obtained by calculating the outer-product matrices of the dictionary elements in the columns of *D*. The state time course was calculated using the mixing matrix *M*. The shape of the final dFC matrix was 1200×*ROI*×*ROI*. The only hyperparameter of this method is the number of FC states.

### Analytical Flexibility Assessment

We used Python to implement all seven methods and estimate the dFC matrix for each of the 395 preprocessed fMRI subjects’ data, resulting in a total of 2765 dFC matrices. Figure2 shows representative dFC matrices obtained by different methods using the BOLD data of one subject. To ensure comparability, as sliding window based methods (SW, SWC, and DHMM) downsample the time samples, the dFC matrices of other methods were uniformly downsampled prior to further analysis to match the resolution of the sliding window based methods. The output of each method was converted to a common format of a 38×96×96 dFC matrix, resulting in 7 dFC matrices of the same size per subject. This resulted in a dFC results array of dimensions *dFC*(*subject, method, time, ROI, ROI*)*∈* (395, 7, 38, 96, 96). The dFC results array was also reshaped in some cases to *dFC*(*subject, method, time, functionalConnection*)*∈* (395, 7, 38, 4560) by vectorizing the lower triangle of 2D FC matrices of shape *ROI*×*ROI* into *functionalConnection*×1. To assess the analytical flexibility of the dFC results across methods, we calculated the 7×(7−1)/2 pairwise similarity using several similarity metrics, including Spearman correlation, Pearson correlation, Euclidean distance, and mutual information. We evaluated the similarity between each pair of methods in terms of correlation of their: dFC matrices *dFC*(*subject, method*, :, :, :) (overall similarity), time courses of functional connections *dFC*(*subject, method*, :, *ROI, ROI*) (temporal similarity), FC patterns at each time point *dFC*(*subject, method, time*, :, :) (spatial similarity), and graph properties such as degree and clustering coefficient (see Figure27 and Figure28 in Supplementary material). The equations used to calculate each of these similarity metrics are as follows:

**Fig. 2.**
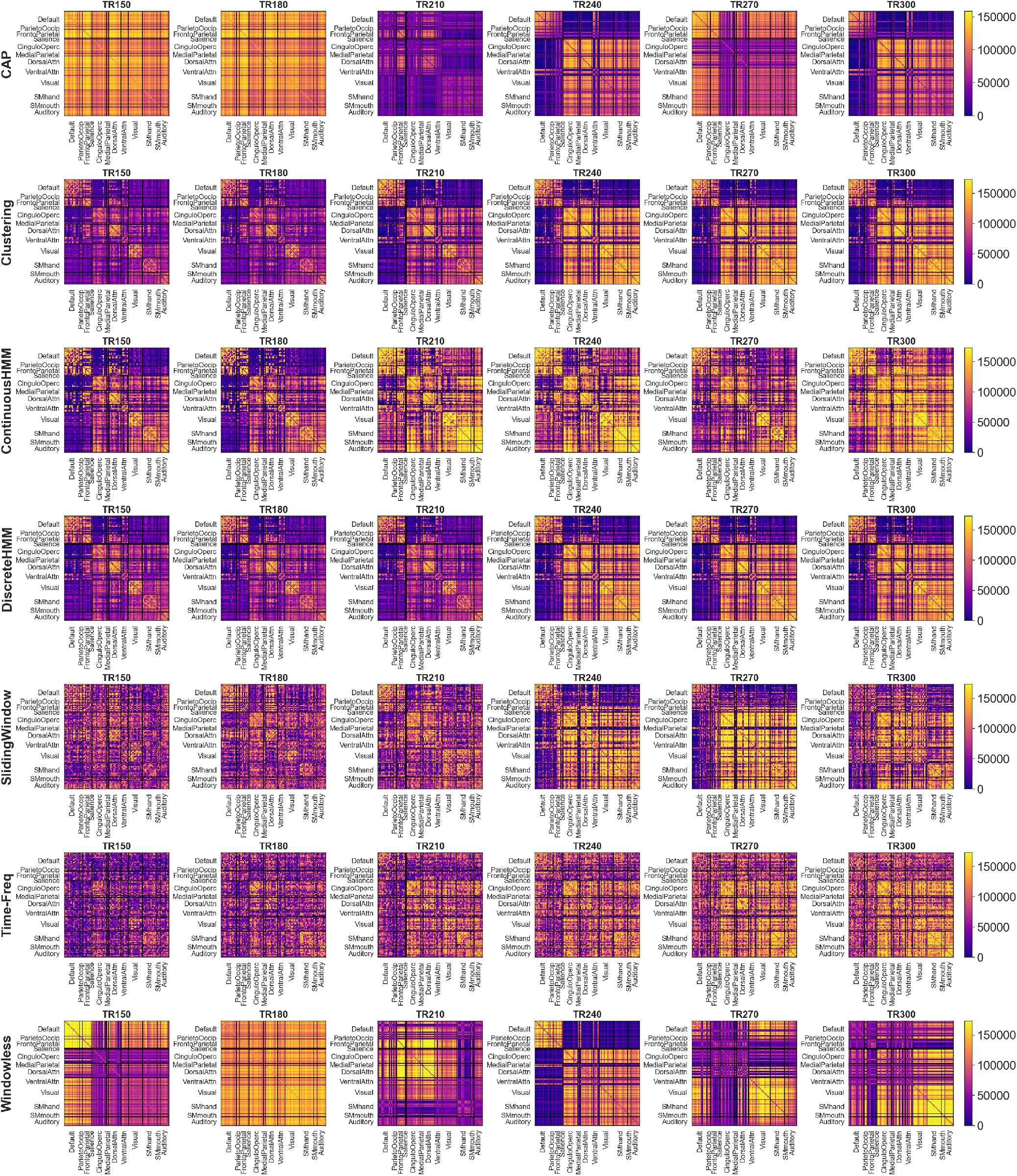
A sample segment of dFC matrices of one subject obtained by the implemented methods: each row shows the dFC matrix obtained by each method and each column shows the corresponding time point, or TR. Each FC matrix has a size of *ROI × ROI* and ROIs are divided into 12 RSNs. To account for the different value distributions of each method and better comparability, the dFC matrices were normalized by ranking the values, similar to the approach used when calculating Spearman correlation.

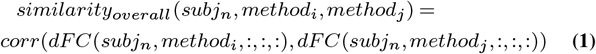

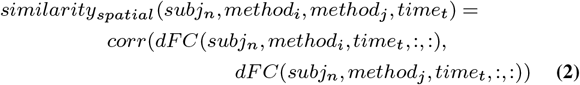

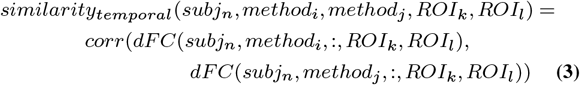

Additionally, we compared the correspondence between the inter-subject correlations of different methods to obtain the inter-subject similarity:

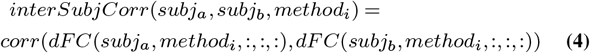

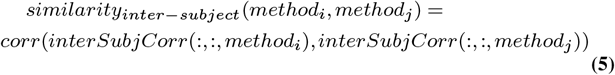

To identify groups of methods that produce similar results, we used hierarchical clustering with the Ward method to analyze the similarity values calculated using the afore-mentioned metrics (Spearman correlation, Pearson correlation, Euclidean distance, and mutual information). For correlation-based metrics, 1 − *correlation* was used as the distance between the methods. The hierarchical clustering allowed us to summarize the measured similarity matrices and, moreover, group together methods that exhibited a higher degree of similarity.

## Results

### DFC assessment methods can be grouped into 3 categories based on the similarity of their results

Figure3a shows the overall similarity obtained by Spearman correlation of the subject-specific dFC matrices computed using the seven selected methods (an illustration of sample dFC matrices obtained by each method can be seen in Figure2). The correlation values were averaged over subjects and repeated for each session, as indicated in equation1 (for results obtained by other metrics see Supplementary material: Pearson correlation, Figure24; Euclidean distance, Figure25; and mutual information, Figure26; see Figure22 in Supplementary material for the results of two-way ANOVA test on the effect of session (day and direction) on the overall similarity values). Correlation values show a range between weak to strong similarity between dFC methods. The average Spearman similarity of all pairs over all subjects was 0.38, indicating a moderate degree of similarity. However, the corresponding standard deviation was high (0.18; variance: 0.032), about half of the average similarity value. This variability was also significantly larger than the average variance of pairwise similarities over subjects (SD: 0.076, variance: 0.0058). In other words, dFC similarity variability over method pairs was five times higher than over subjects.

**Fig. 3.**
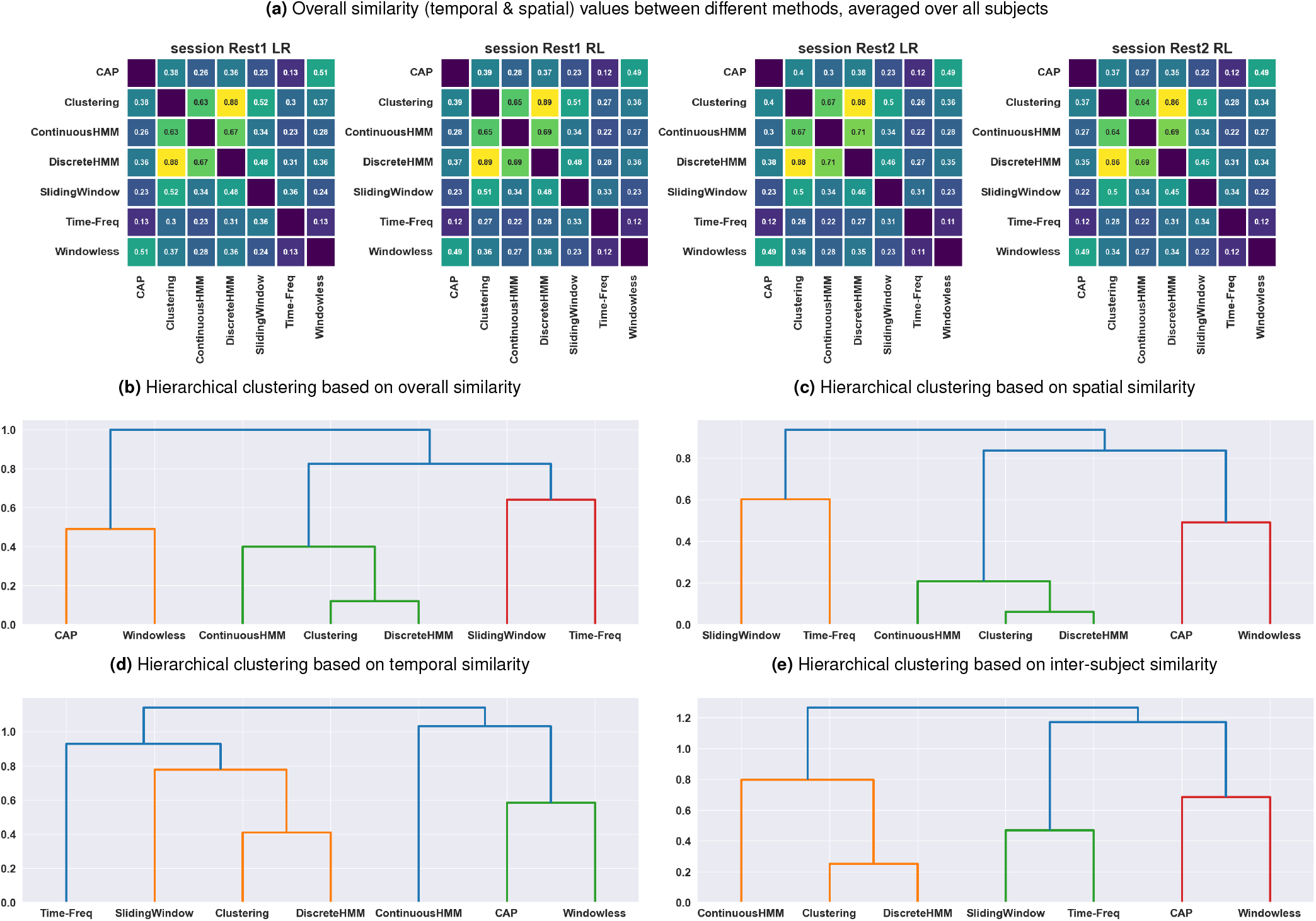
Similarity between dFC patterns obtained from seven methods as assessed by Spearman correlation: (a) The overall (combined spatial and temporal) similarity values averaged over subjects show considerable variation over different method pairs in all sessions; hierarchical clustering based on average similarity (b) overall (comparing dFC matrices), (c) spatial (comparing FC matrices at each time point), (d) temporal (comparing time courses of each functional connection), and (e) inter-subject (comparing intersubject correlation values) measured by Spearman correlation. The hierarchical clustering results suggest that the methods can be grouped into three groups: 1) Clustering, Continuous HMM, and Discrete HMM; 2) CAP and Window-less; 3) Sliding Window and Time-Frequency. A slightly different pattern was obtained through temporal similarity assessment, which is due to decreased similarity of Continuous HMM and Time-Frequency with the methods in their group. The general similarity between the methods was relatively low, particularly for inter-group comparisons. For results obtained by other metrics see Supplementary material: Pearson correlation, Figure24; Euclidean distance, Figure25; and mutual information, Figure26

We used hierarchical clustering analysis on the pair-wise similarity matrices to investigate and summarize variation between dFC methods. The hierarchical clustering based on these average correlation matrices (Figure3b) suggests that the methods can be classified into three groups: 1) Clustering, Continuous HMM, and Discrete HMM; 2) CAP and Window-less; 3) Sliding Window and Time-Frequency. Although the definition of these groups highly depends on the cutoff value of the clustering, we observed high intra-group similarities and lower inter-group similarities for these 3 groups. Most subjects had the same hierarchical clustering as the one observed on the average across subjects (Figure21 in Supplementary material).

The distribution of these similarity patterns over subjects (Figure20 in Supplementary material) indicates that the similarity between certain pairs of methods, notably those involving Time-Frequency, displays greater variability over subjects. This observation suggests that the observed similarity between these methods is more likely influenced by individual differences in BOLD signals than mathematical similarity of their analyses. In other words, the variation in similarity across subjects implies that individual-specific characteristics may play an important role in the similarity between these methods. However, for other pairs of methods, more consistent similarity profiles were observed over subjects.

### Similarity in spatial patterns of dFC results follows the overall similarity patterns

The spatial similarity between the dFC matrices obtained from different methods was estimated using Spearman correlation between their corresponding spatial patterns (FC matrices) at each time point for each subject (see equation2). The resulting spatial similarity was then averaged over both time and subjects. Hierarchical clustering analysis of the average spatial similarity matrix (see Figure3c) revealed the same hierarchical structure as the one revealed by average overall similarity. The clustering analysis identified the same three groups, suggesting that their spatial similarities follow the same intra- and inter-group pattern.

### Across methods dFC assessments reveal lower temporal similarity compared to spatial similarity

The temporal similarity between dFC matrices of each pair of methods was assessed using Spearman correlation between the time courses obtained by the methods for each functional connection (see equation3). The resulting temporal similarity matrices were then averaged over functional connections (*ROI*×*ROI*) and subjects. While two methods may produce dFC matrices that have similar spatial patterns, the corresponding temporal patterns may differ. Across methods, spatial similarity focuses on the spatial patterns and values of functional connections at a given time, while temporal similarity focuses on the consistency of the time evolution of dFC for each functional connection. The hierarchical clustering based on the average over subjects temporal similarity matrix (Figure3d) reveals an overall decrease in similarity between methods compared to spatial (and overall) similarity. This was particularly noticeable for the similarities between CHMM and TF with other methods in their groups, which yielded considerably lower temporal similarity compared to the corresponding spatial similarity. This suggests that although CHMM and TF exhibit similar spatial patterns to the methods in their groups, they do not exhibit equally similar temporal dynamics (see Figure31 and Figure32 in Supplementary material).

### Inter-subject similarities reflect the previously observed overall similarity patterns

To assess an intersubject similarity between methods, dFC matrices estimated by each method were used to measure inter-subject correlations (395×(395−1) / 2 values) (see equation4). The inter-subject values obtained for each method were then compared and the similarity between methods was measured using Spearman correlation (see equation5). The resulting correlation values (Figure3e), indicating inter-subject correlation correspondence between methods, revealed the same three groups of methods in terms of their similarity. This implies that if one method identifies two subjects as similar based on their dFC patterns, it is more likely that another method from the same group will also identify those two subjects as similar. This finding suggests that the methods grouped together not only demonstrated similarity in capturing subject-level patterns but also exhibited consistency in capturing inter-subject patterns. For results obtained using Fractional Occupancy (FO) as the feature used for assessing the inter-subject correlations see Figure29 in Supplementary material.

### The variability over method is comparable to the variability over time for most functional connections

The variance of dFC,*dFC*(*subject, method, time, functionalConnection*), was calculated over time and method, across functional connections and subjects. To account for the different value distributions of each method, the dFC matrices for each subject, *dFC*(*subject*_*n*_, *method*_*i*_, *time, functionalConnection*),were normalized by ranking the values, similar to the approach used when calculating Spearman correlation. We calculated variance over time, *var*(*dF C*(*subject, method*, :, *functionalConnection*)),averaging over method, and the variance over method,*var*(*dF C*(*subject*, :, *time, functionalConnection*)), averaging over time. Therefore, both calculations resulted in an array of *subject* × *functionalConnection* values (one ‘temporal variance’ and one ‘methods variance’ value per functional connection and per subject). The results, shown in Figure4a, indicate that the variation over dFC methods was comparable to the variation over time for most functional connections and subjects, leading to an average ratio of *var*_*method*_*/var*_*time*_ = 0.95, or an average ratio of *SD*_*method*_*/SD*_*time*_ = 0.97. For the variability of the results obtained across method pairs and method groups see Figure37 and Figure38 in Supplementary material. Moreover, Figure4b highlights the pair of RSNs with functional connections that exhibited higher variability over method than over time. These functional connections mainly include connections between the Default Mode and other networks, such as Parieto-Occipital, FrontoParietal, Salience, Cingulo-Opercular, MedialParietal, Dorsal Attention, and Ventral Attention Networks, as well as most intra-network connections, with the exception of the Cingulo-Opercular Network.

**Fig. 4.**
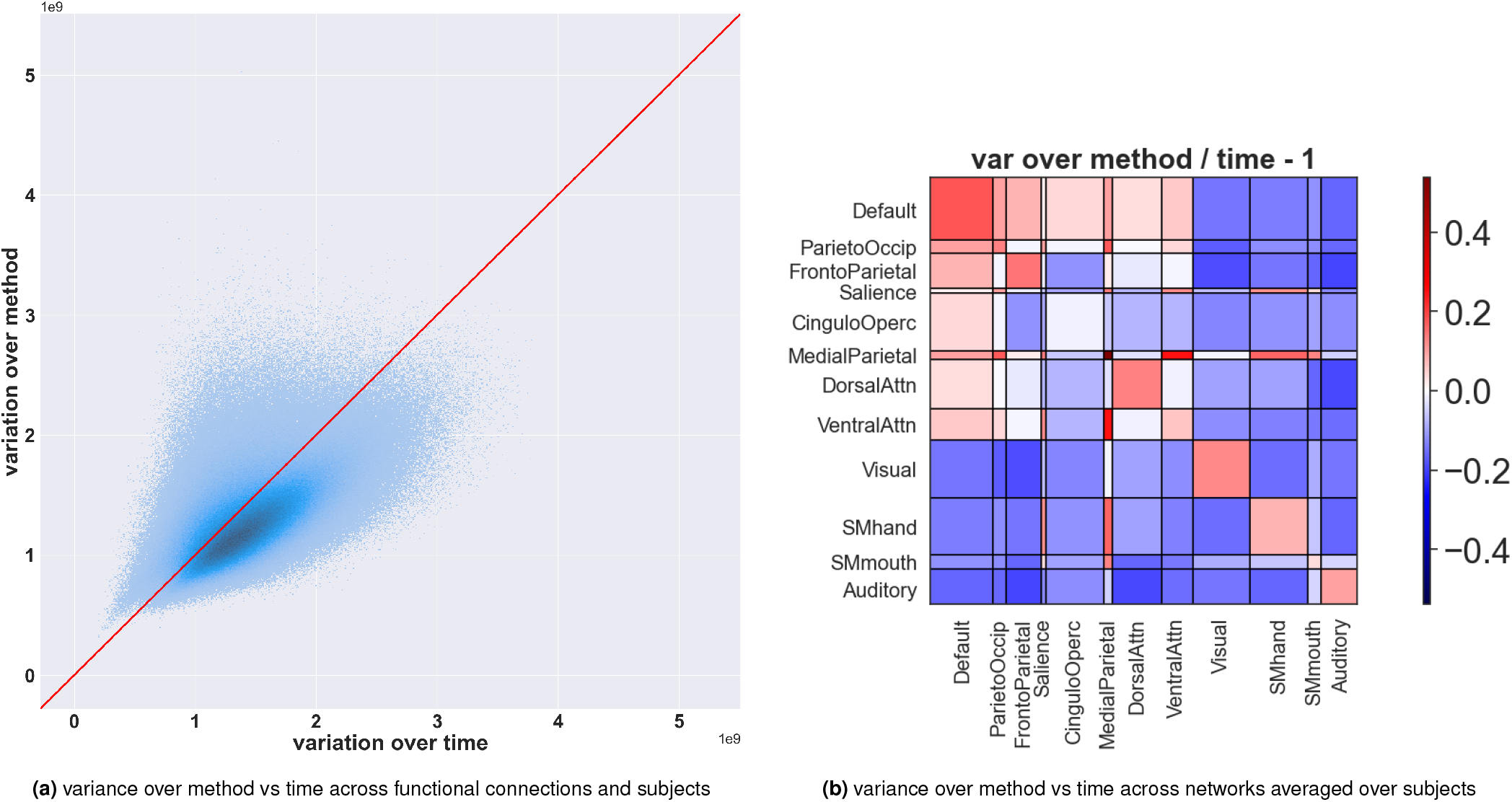
a) A scatter plot of the variance over methods versus the variance over time for each functional connection and each subject (*subj*× (*ROI*× (*ROI* −1)/2) points in total) indicates that the variance over dFC methods is comparable to the variance over time, with an average ratio of *var*_*method*_*/var*_*time*_ = 0.95 (*SD*_*method*_*/SD*_*time*_ = 0.97). b) Ratios of variance over method divided by variance over time across functional connections averaged over subjects. The ratios were subtracted by one (*ratio −* 1) so that zero values (white) correspond to equal variance over method and time. The current figure highlights Resting-State Networks (RSNs) pairs with functional connections that are more variable over method than time (red) and those more variable over time than method (blue). The variance values corresponding to functional connections belonging to each pair of RSNs were averaged to yield a single variance value prior to computing the ratios. The plot shows that the functional connections between RSNs such as the FrontoParietal and Ventral Attention networks exhibited equal variance values over time and method, while the functional connections between RSNs such as the Default Mode and Parieto-occipital, and Default Mode and FrontoParietal Networks, and most intra-network connections exhibited higher variation over method than over time.

**Fig. 5.**
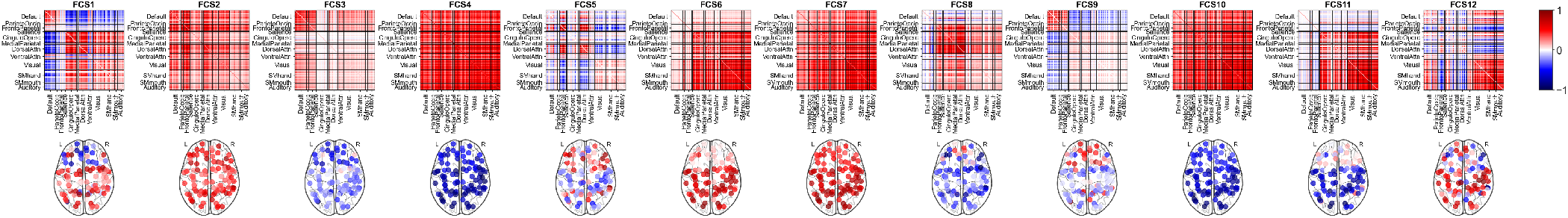
state FC matrices of the CAP method. Among the 12 state FC matrices, FCS4 is likely to be associated with artifacts.

**Fig. 6.**
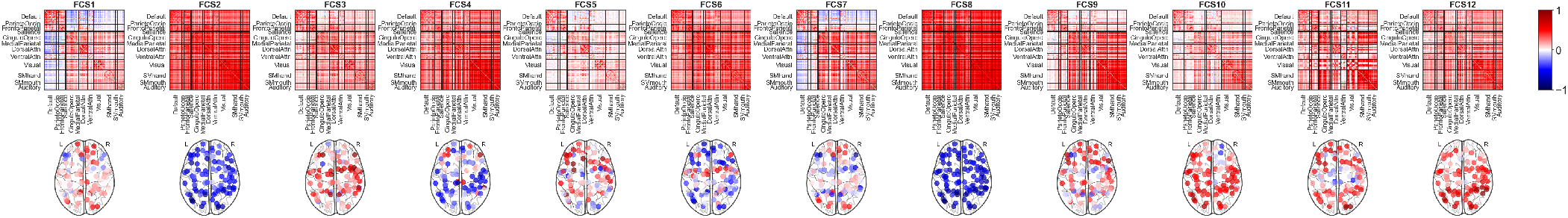
state FC matrices of the Clustering method. Among the 12 state FC matrices, FCS8 is likely to be associated with artifacts.

**Fig. 7.**
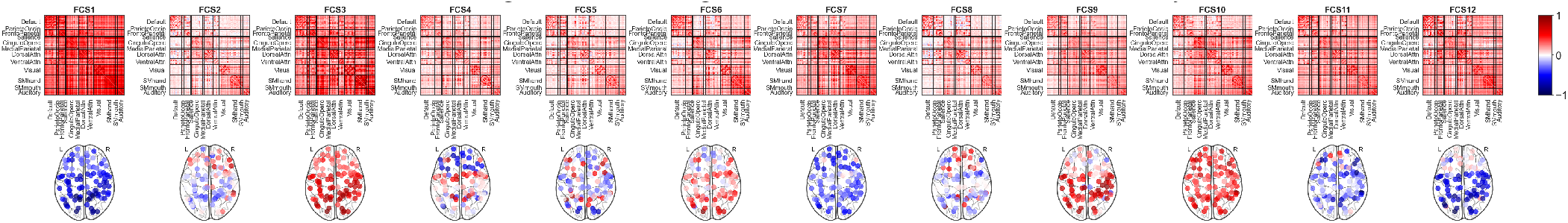
state FC matrices of the Continuous HMM method.

**Fig. 8.**
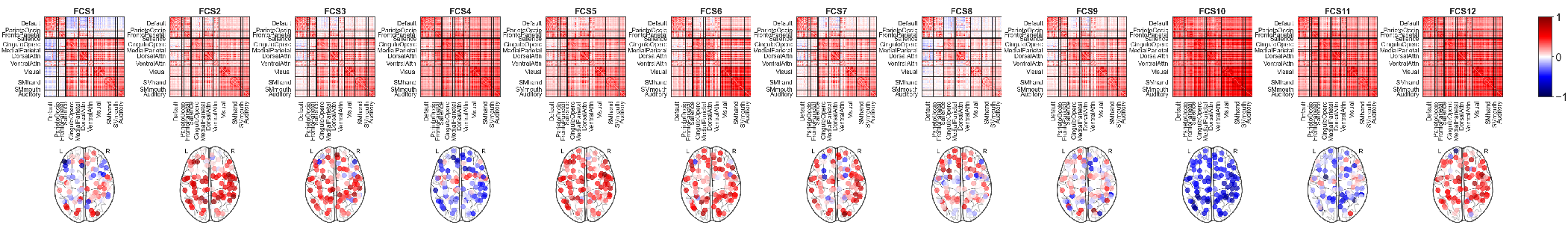
state FC matrices of the Discrete HMM method.

**Fig. 9.**
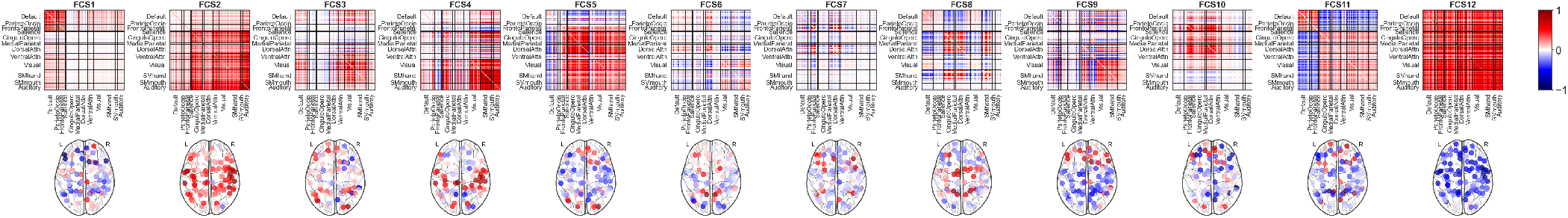
state FC matrices of the Window-less method. Among the 12 state FC matrices, FCS12 is likely to be associated with artifacts.

**Fig. 10.**
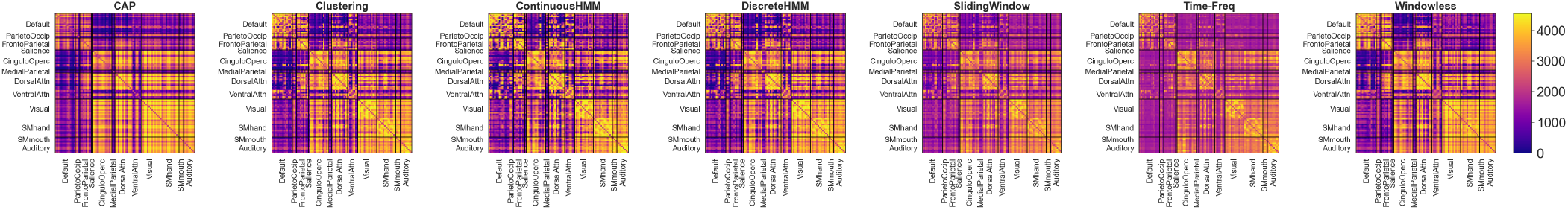
The average dFC, or static FC, for each method was calculated by averaging the dFC matrices over time and subjects, and then normalizing the values by ranking them to make the results more comparable. While the overall pattern of the static FC matrices was similar across methods, there was some degree of dissimilarity observed between the matrices produced by TF compared to the other methods.

**Fig. 11.**
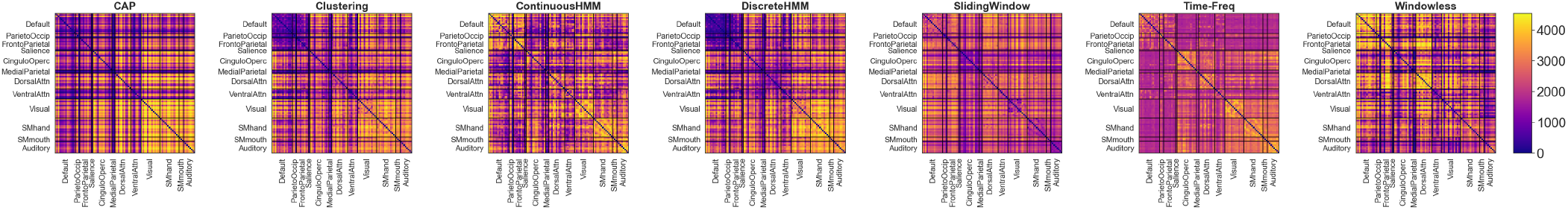
The temporal variability of dFC matrices evaluated for each method by computing the variance of the dFC matrices over time, averaged over subjects: The results were rank-normalized to make them more comparable. The resulting temporal variance matrices revealed significant differences between methods. This can be observed by comparing the high-variance and low-variance functional connections yielded by each method, indicating that functional connections that are considered dynamic by one method may not be considered as dynamic by another method.

**Fig. 12.**
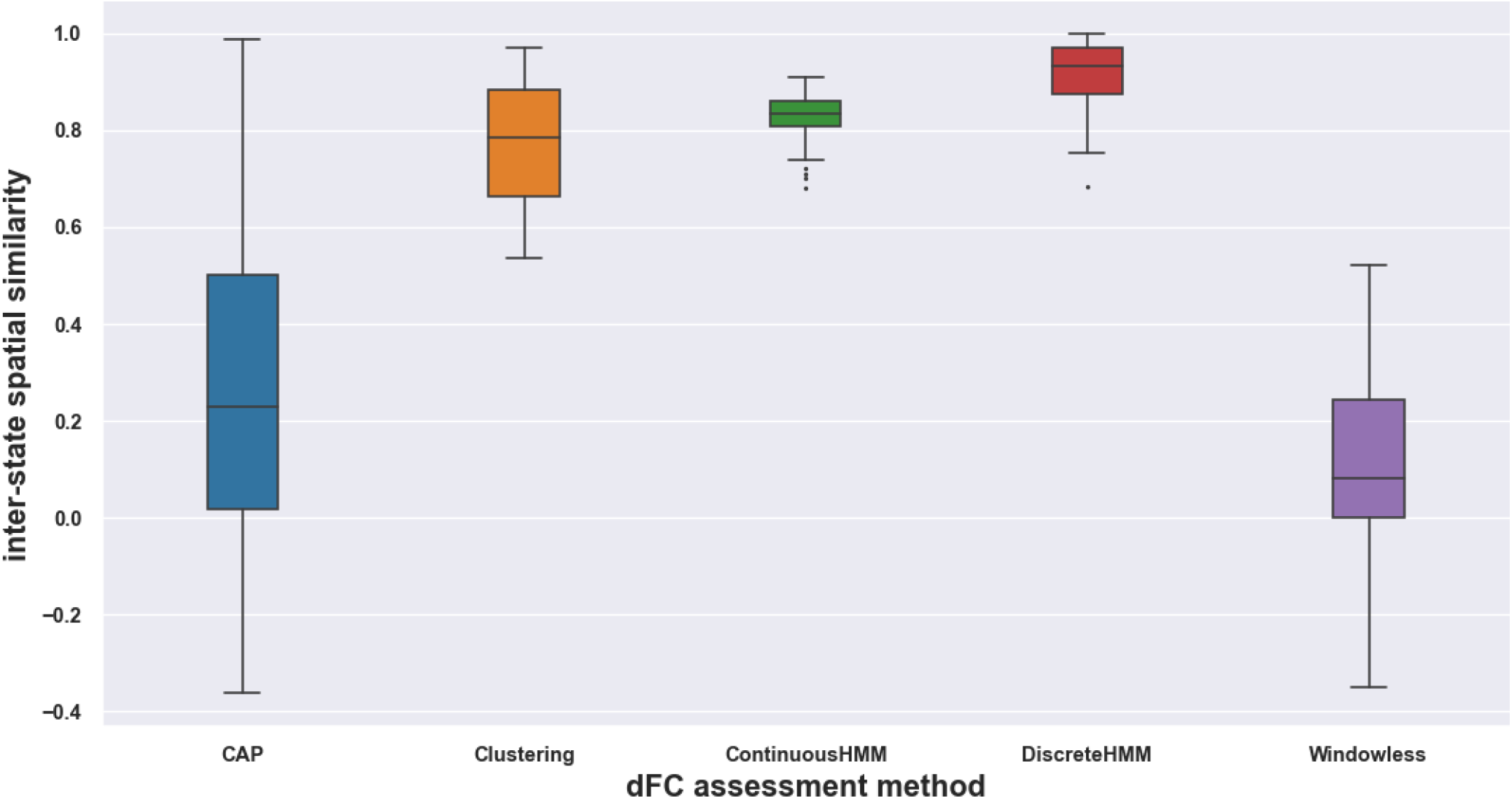
The figure shows the similarity distribution of the spatial patterns of states across different methods. The results indicate that SWC, CHMM, and DHMM had a more consistent range of similarities, with higher average similarity between states in terms of their spatial patterns. In contrast, CAP and WL exhibited a wider range of similarities and lower average similarity between states.

**Fig. 13.**
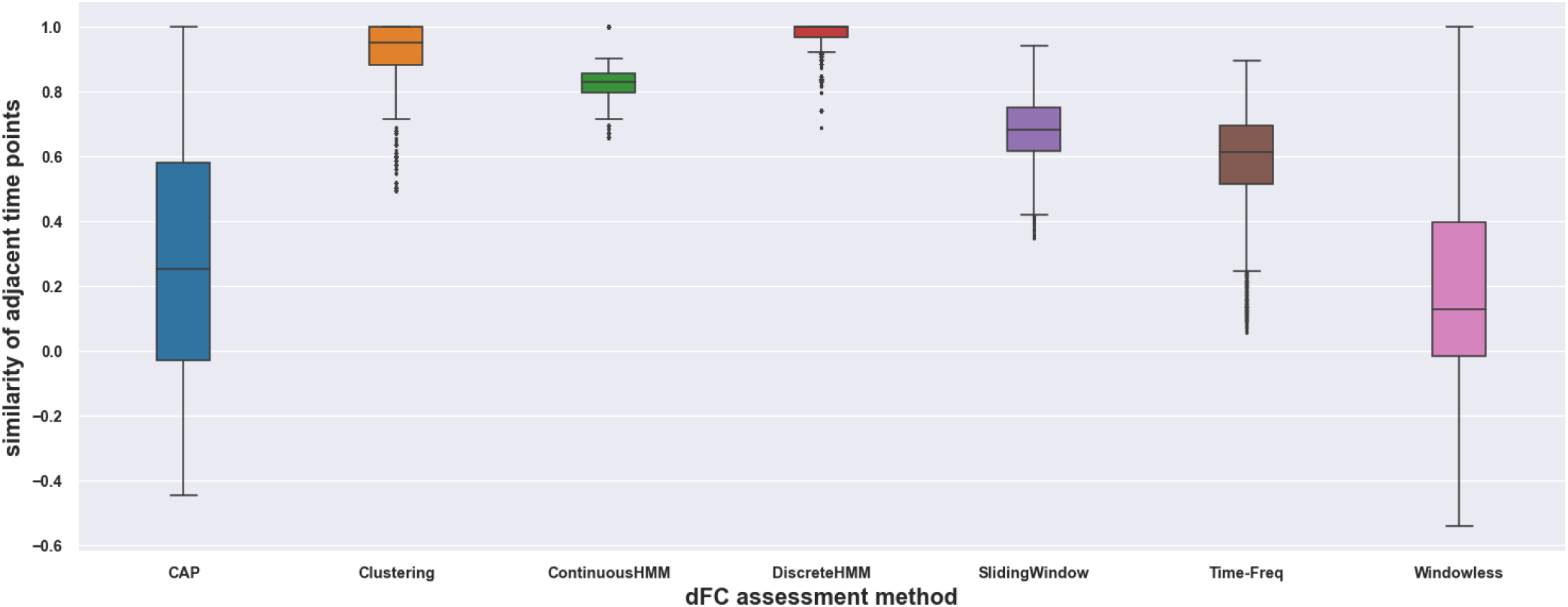
Similarity of adjacent time points, or smoothness, measured by finding the Spearman correlation of each FC matrix and its subsequent FC matrix in time and averaged over all pairs of adjacent FC matrices and averaged over subjects. Higher average similarity implies higher smoothness in the dFC matrices obtained by a specific method. The results show that methods such as SWC, CHMM, and DHMM demonstrate higher smoothness, likely due to their locality of neighboring time points assumption. Conversely, methods such as CAP and WL exhibit lower smoothness, likely because they do not assume locality of neighboring time points.

**Fig. 14.**
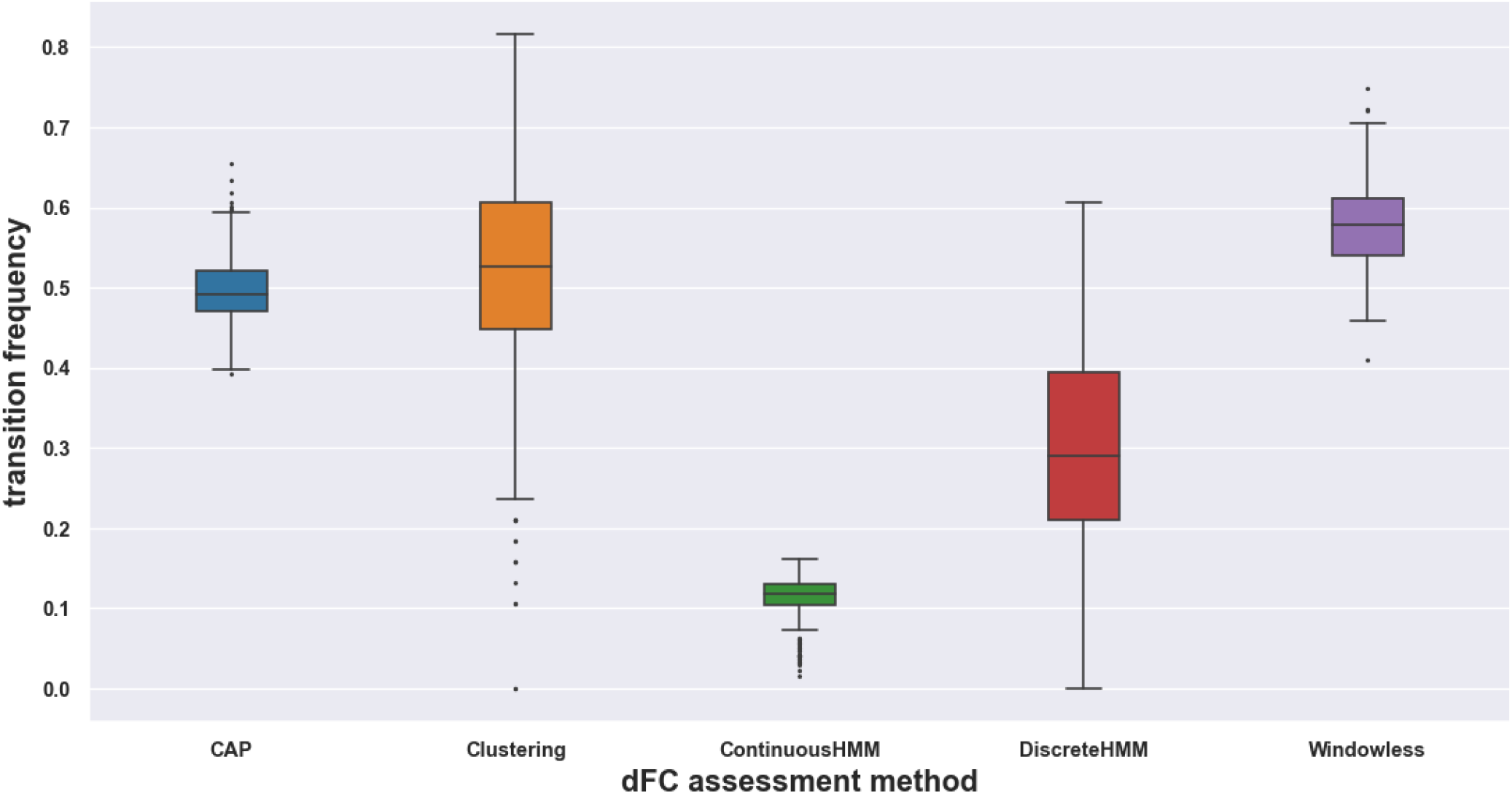
The frequency of state transitions for each state-based method was calculated by dividing the number of state transitions by the total number of time samples in the dFC matrix. Normalization was applied to account for differences in time resolution among methods. The results indicate that methods such as CHMM and DHMM have a lower frequency of state transitions, likely due to their assumption of smooth state transitions and lack of instantaneous reconfigurations. Conversely, methods such as CAP, SWC, and WL have a higher frequency of state transitions, likely reflecting the presence of instantaneous reconfigurations in their dFC matrices.

**Fig. 15.**
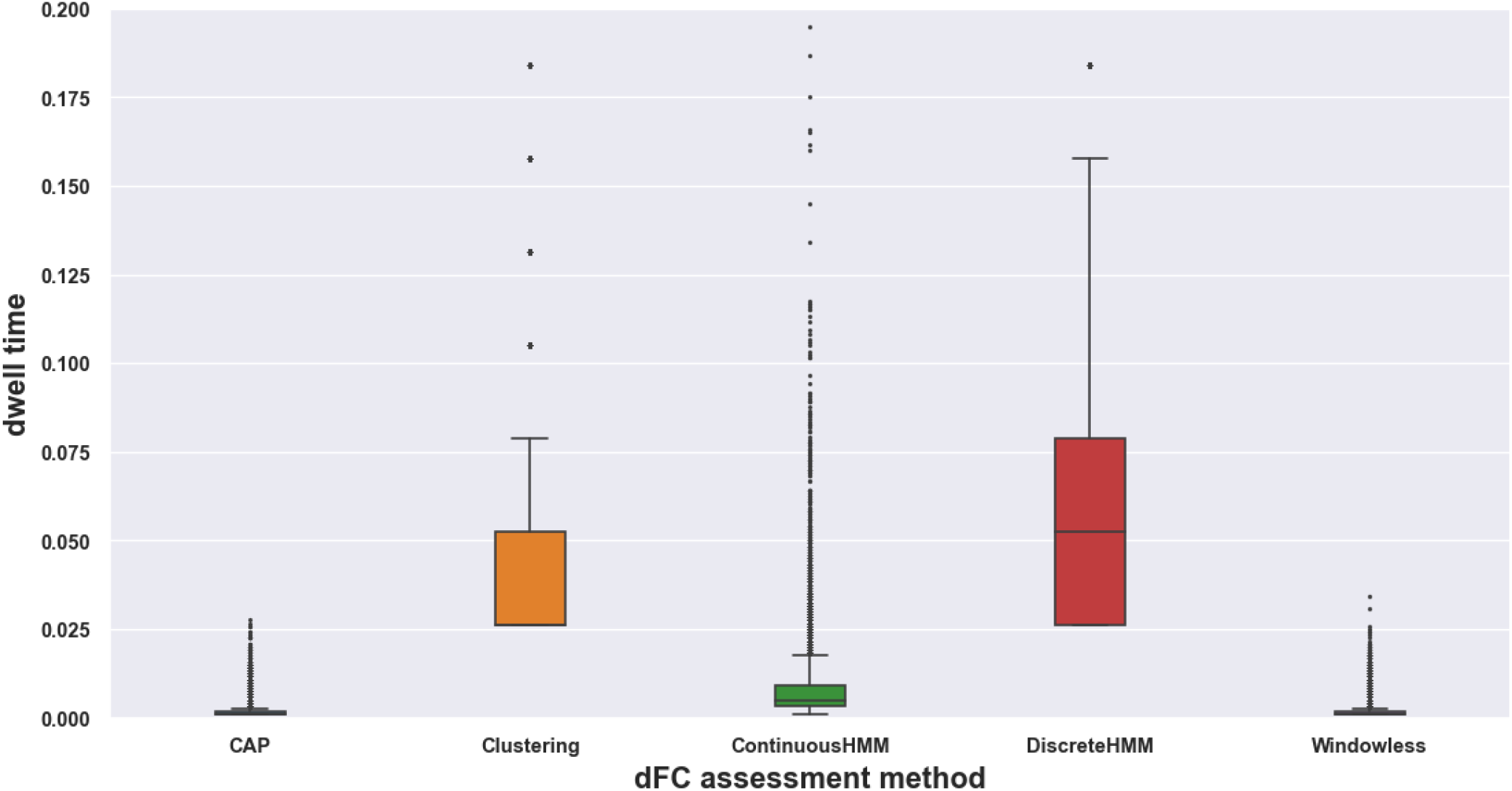
Distribution of dwell times, defined as the time elapsed between state transitions, across different state-based methods. The dwell times were normalized by the total number of time points of each subject. Our analysis reveals that CAP and WL had significantly shorter dwell times compared to the other three methods. On the other hand, DHMM had the longest dwell times, up to 0.2 (or 240 TR equal 172.8 seconds). Interestingly, SWC exhibited relatively long dwell times despite having less temporal dependence than HMM-based methods.

**Fig. 16.**
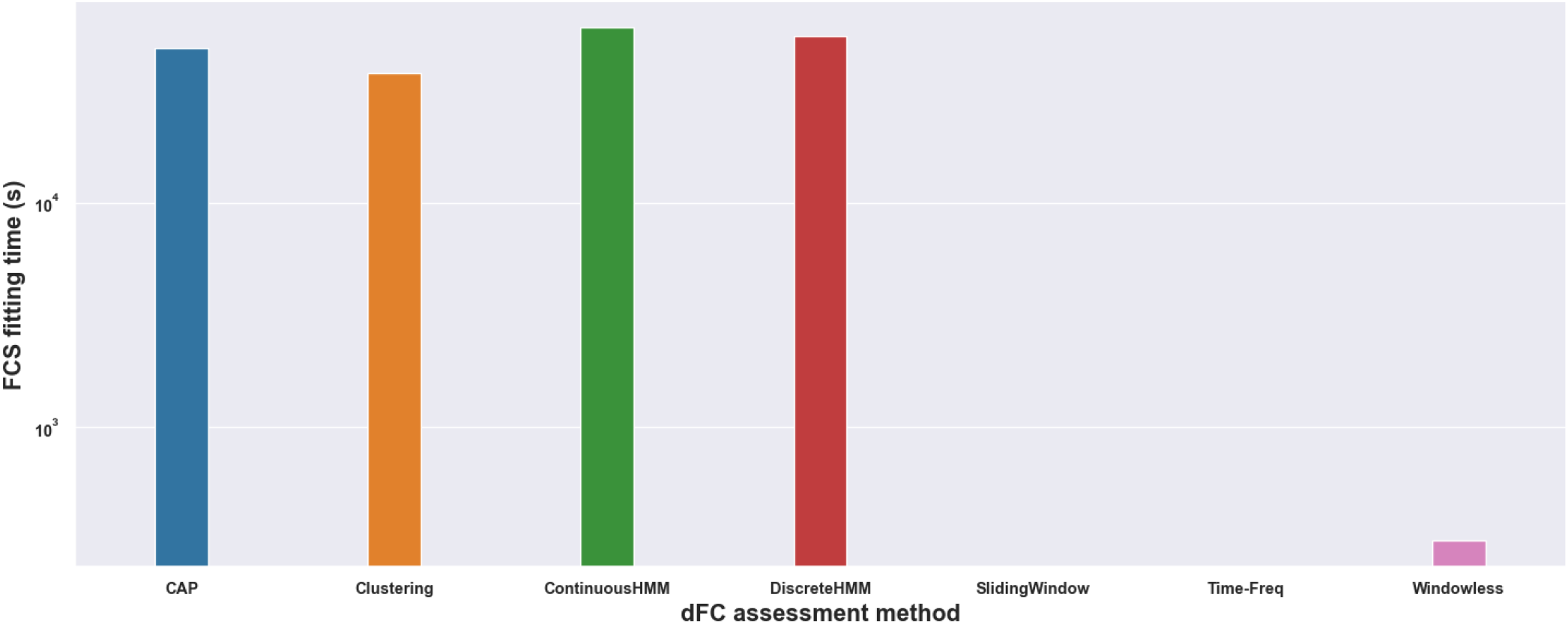
The computation time for fitting states of each state-based method on data of all subjects. The bar chart shows that CHMM has the longest computation time and WL has the shortest, taking only a few seconds. SW and TF have zero fitting time, since they are state-free methods.

**Fig. 17.**
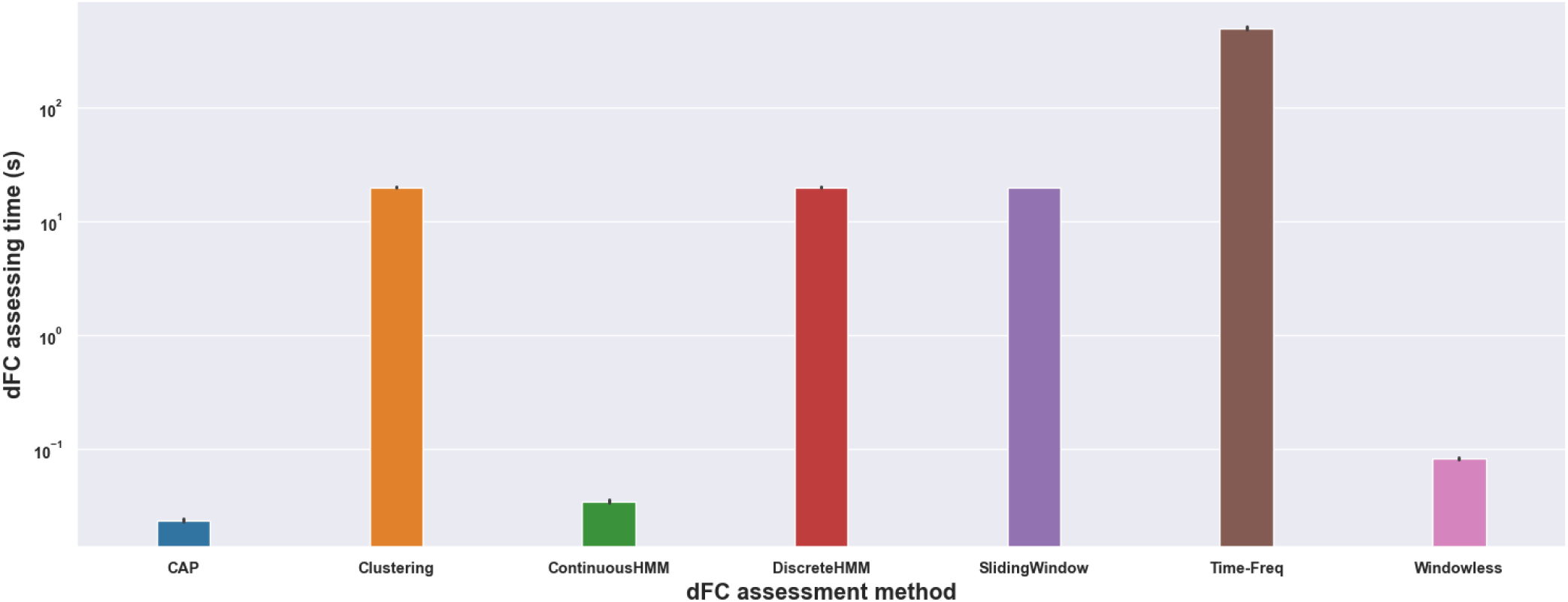
Comparison of average computation time for estimating dynamic functional connectivity (dFC) of a subject using different methods. State-based methods such as CAP, CHMM, and WL have the shortest estimation time, taking only a few seconds. SWC and DHMM still have a relatively long computation time, since their dFC assessment relies on Sliding Window analysis of the data. In contrast, state-free methods have the longest computation time, taking several minutes. TF has the longest computation time, with an order of magnitude higher than the others.

**Fig. 18.**
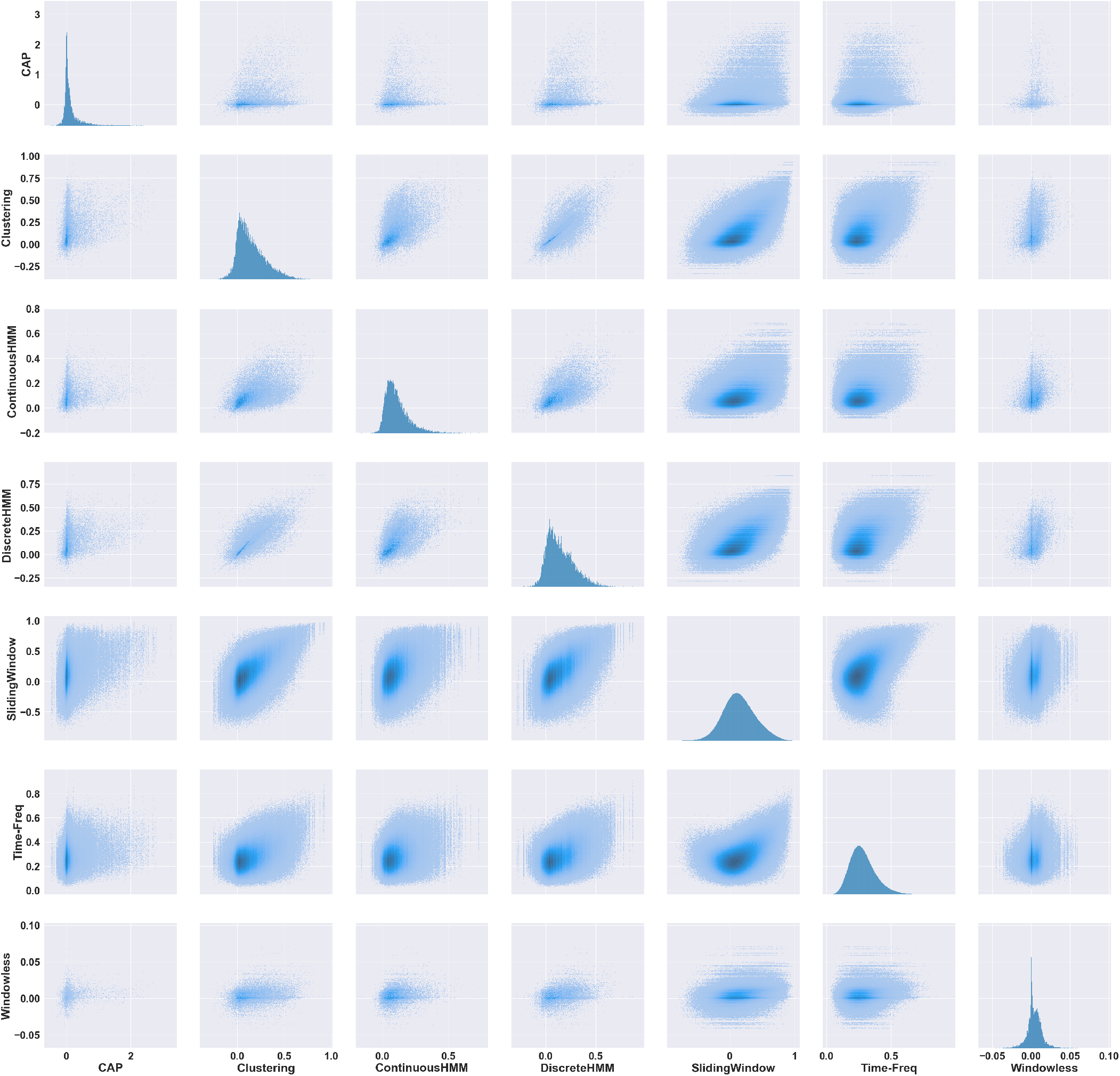
Single and joint distribution of dFC values obtained by different methods. For computation reasons only the first 100 subjects are included in this analysis. Each distribution exhibits a distinct shape, with some methods showing Gaussian distributions (e.g., SW and TF) and others displaying non-Gaussian distributions. WL and CAP have notably narrow distributions compared to other methods. The joint distributions for some method pairs, such as SWC-DHMM and CHMM-DHMM, show higher correlation, indicating more consistent patterns, while pairs like WL-SW or SWC-TF exhibit relatively uncorrelated joint distributions, suggesting greater variability. These findings highlight the importance of considering distributional characteristics when interpreting and comparing dFC results.

**Fig. 19.**
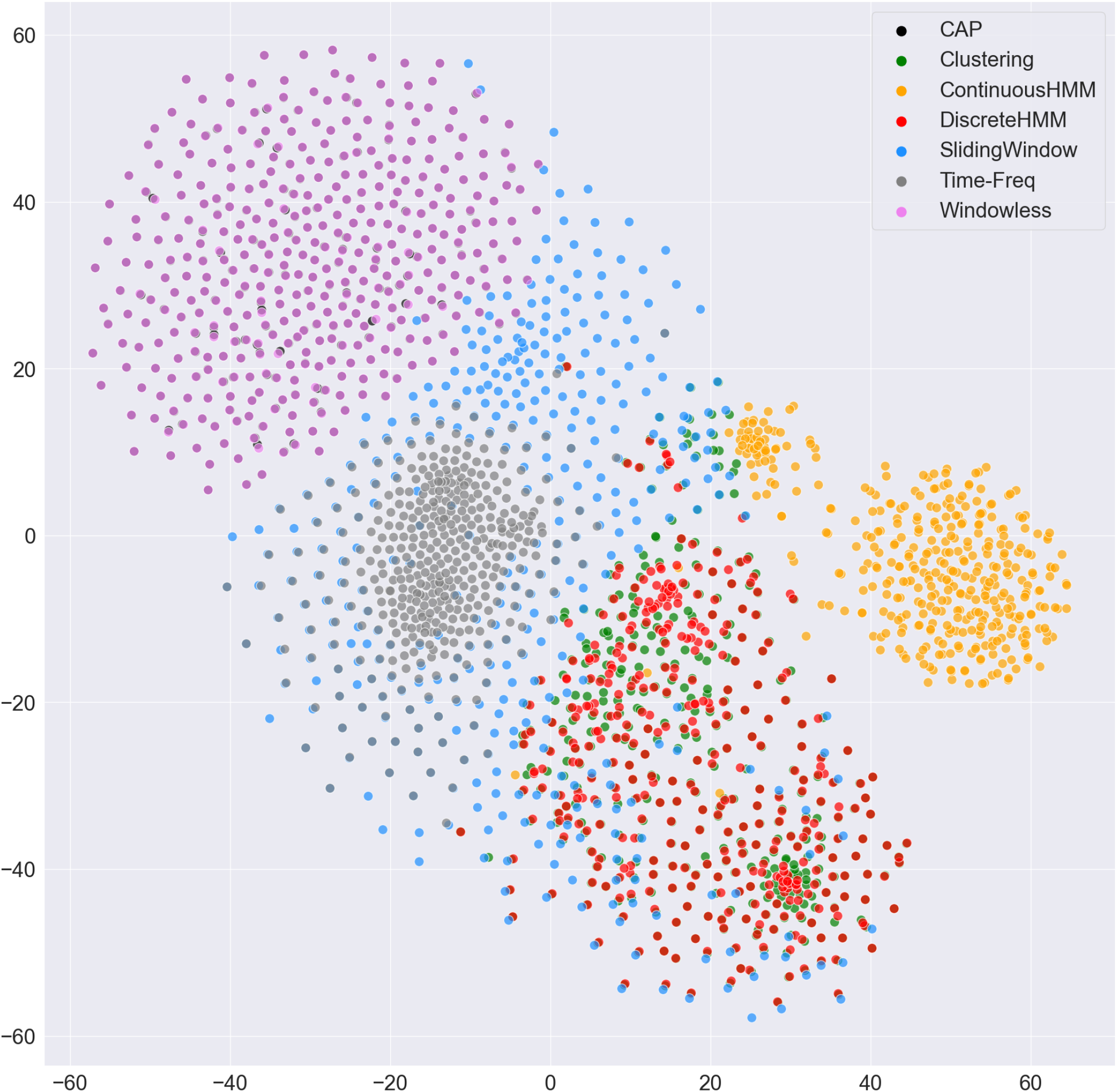
Embedded 2D visualization of subjects’ dFC assessed by different methods. The dimensionality of the dFC matrices was reduced to 1000 dimensions using Principal Component Analysis (PCA), and t-distributed stochastic neighbor embedding (t-SNE) was applied to the transformed data for visualization. Note that the CAP and WL data points overlap. The t-SNE technique facilitates the representation of high-dimensional data in a lower-dimensional space, allowing the visualization of clustering and patterns in the dFC results of subjects across different methods. It is worth noting that due to the dimensionality reduction, a considerable part of the data variation is lost (keeping only 1000 components out of 173,280 in PCA results in approximately 30 percent variance loss). However, the observed patterns in the t-SNE representation align closely with the previous findings. Keeping a different number of components in PCA may result in slightly different patterns.

**Fig. 20.**
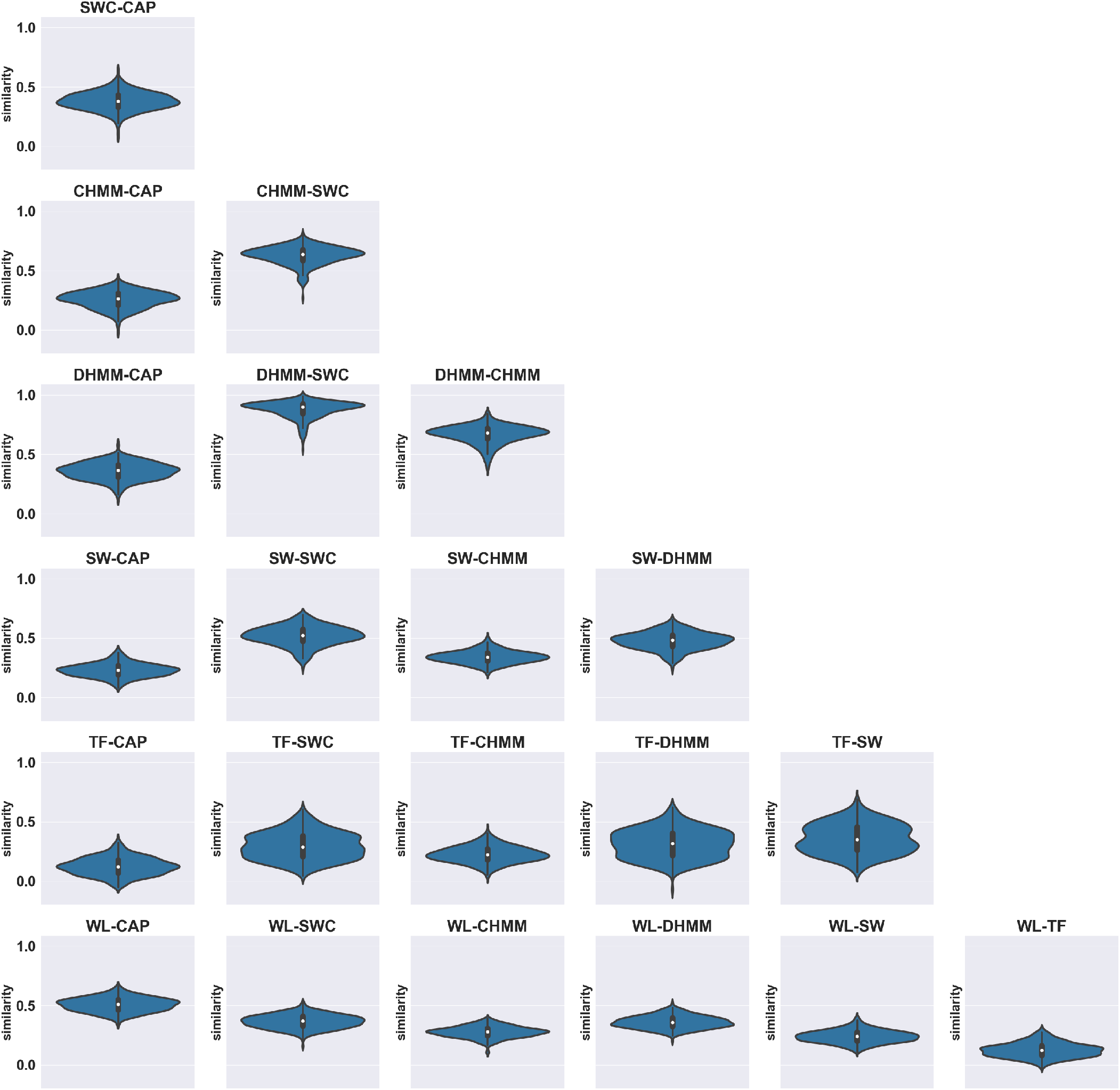
The violin plots display the distribution of overall similarity between dFC matrices for each method pair obtained by Spearman correlation over all subjects. This plot show that the similarity between results of some pair of methods can have considerably diverse values in different subjects, ranging between weak to medium, even relatively strong in some cases. This is mostly the case for the similarity between TF and others. On the other hand, pairs like DHMM-SWC have a high similarity value in all subjects, and pairs such as WL-TF and WL-SW exhibit low similarity values for all subjects. The distributions were estimated using kernel density estimation (KDE) algorithm.

**Fig. 21.**
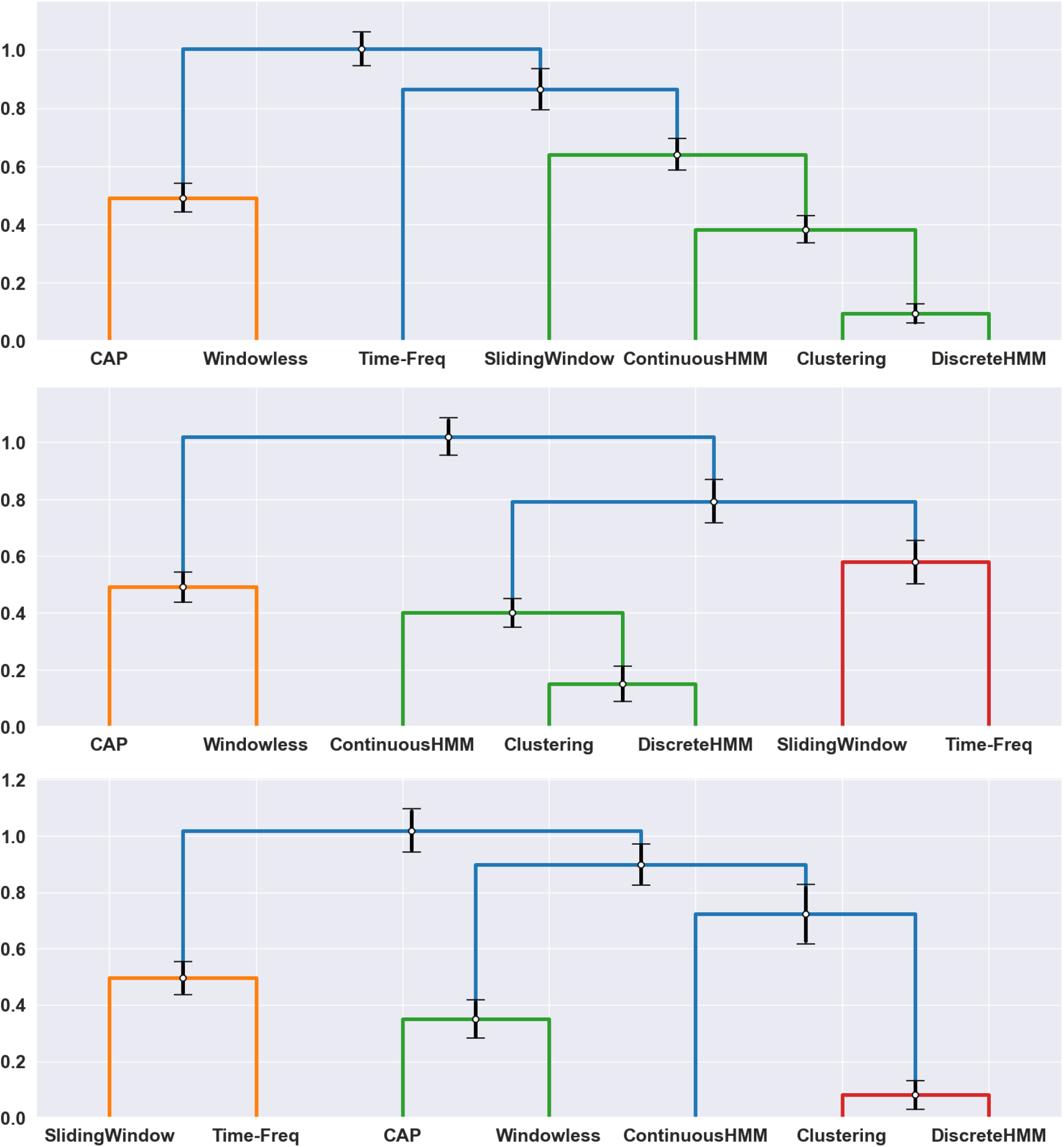
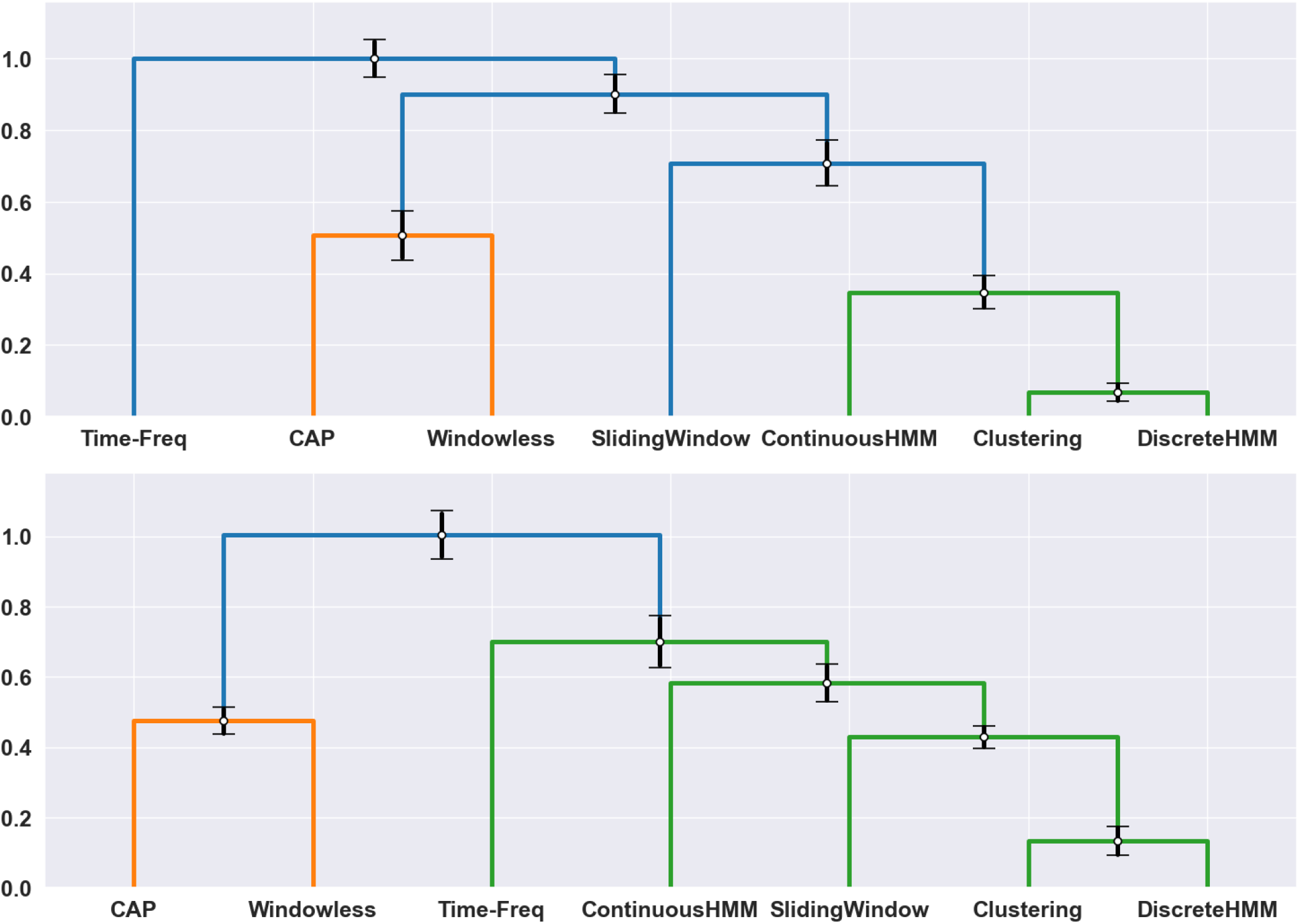
The five most common hierarchical clustering structures across subjects obtained using overall similarity matrix of each subject. For each structure the average and standard deviation interval of the distances are shown on the hierarchical structure. These structures occurred in 110, 144, 21, 66, and 20 subjects (out of 395) respectively. They were selected among a total of 15 existing structures across subjects, as they occurred in at least 10 subjects. The second structure, which was the most common one, is the same structure as the structure obtained using the average overall similarity of all subjects (Figure3b). The variation of structures is mostly due to variation in similarity of Time-Freq, and to a less degree, Continuous HMM with other methods

**Fig. 22.**
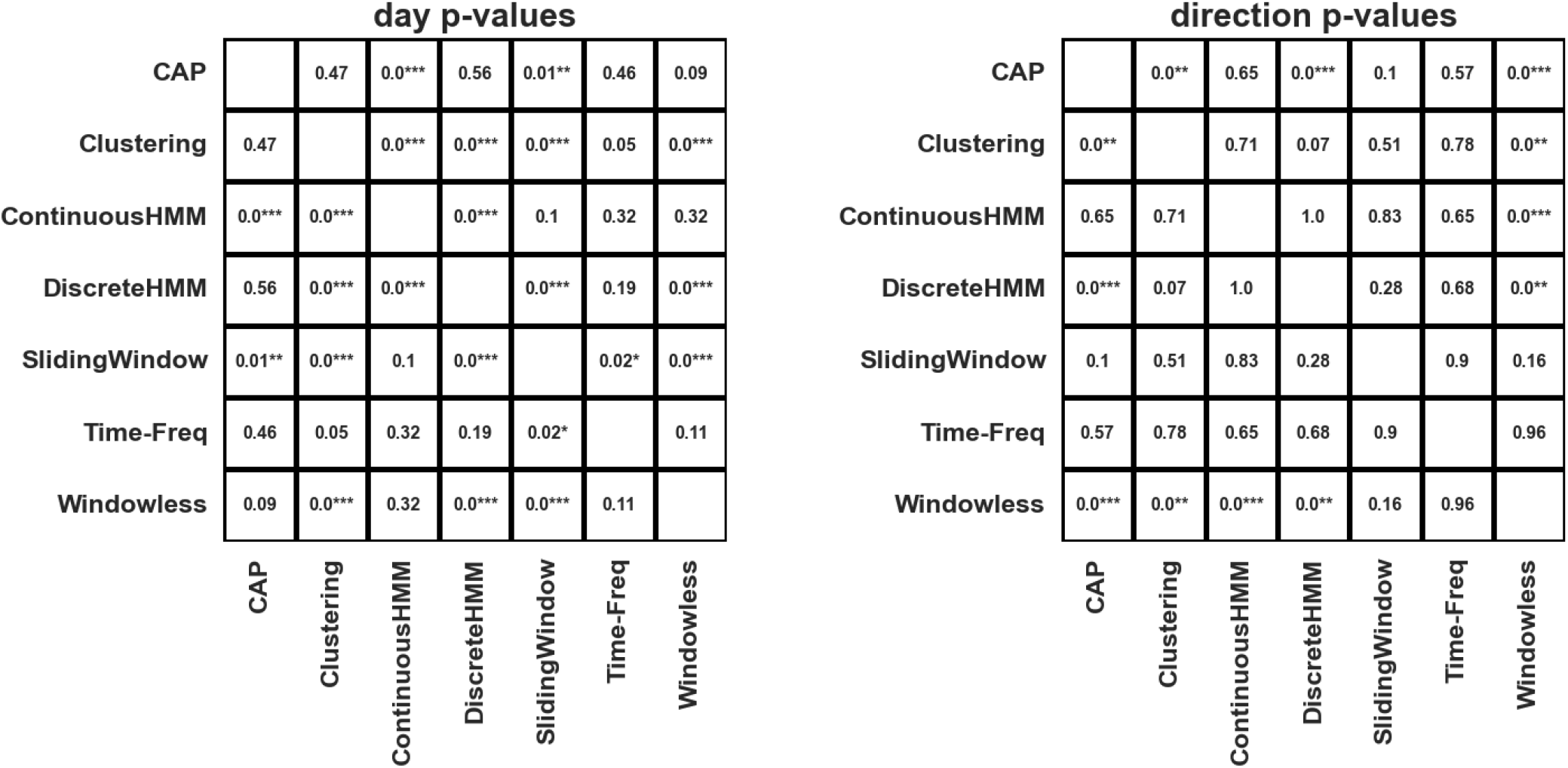
The p-values of the two-way ANOVA on the effect of day (day 1 or day 2) and direction (LR or RL) on the measured similarity of each pair. The similarity of roughly half of the pairs was significantly affected by the day of the scan, while direction had no significant effect on the measured similarities, except for WL and CAP.

**Fig. 23.**
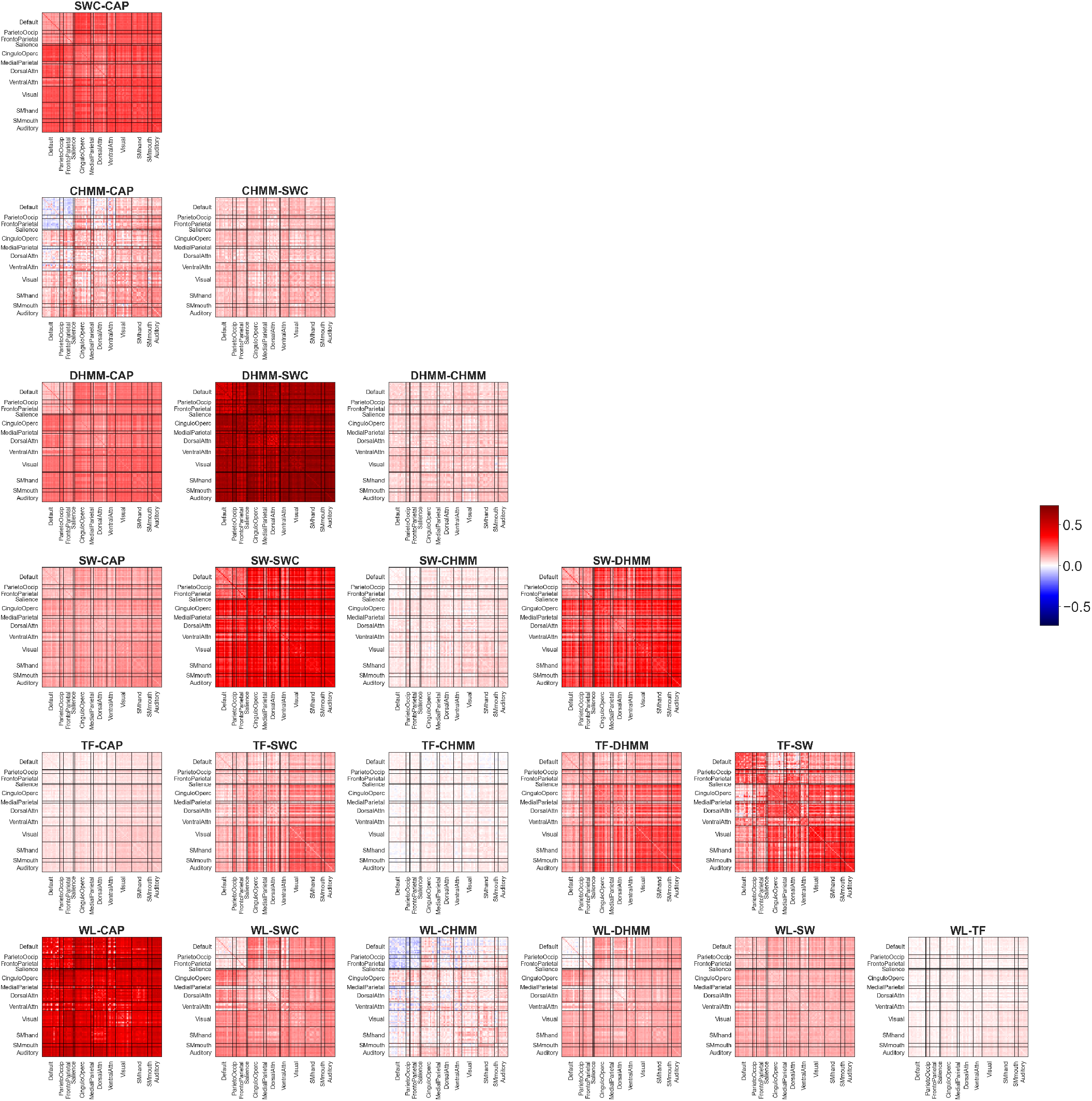
Similarity of dFC matrices obtained by different methods assessed across functional connections using Spearman correlation and averaged over subjects: The resulting similarity matrices highlight areas of high correspondence and low correspondence between the dFC matrices of each method pair, providing a visual representation of the degree of similarity between the methods for each functional connection.

**Fig. 24.**
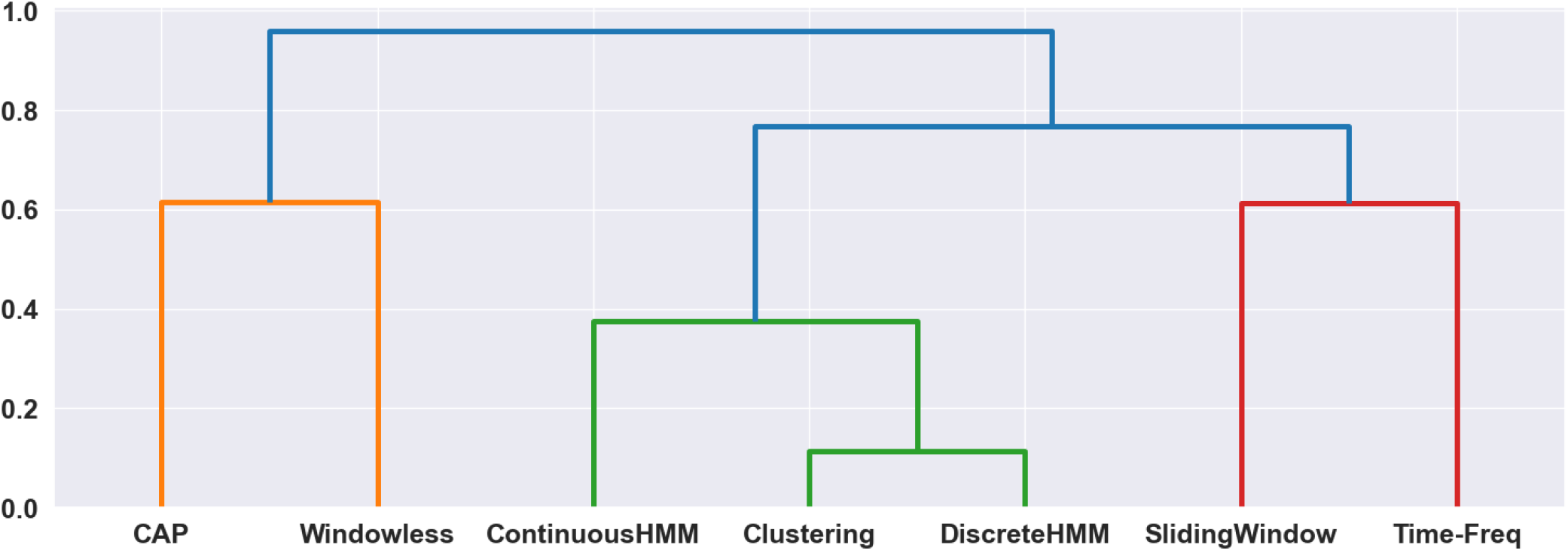
Pearson correlation: Hierarchical clustering of various methods based on the average overall similarity, as determined by Pearson correlation. The results are consistent with those obtained through the use of Spearman correlation, with the same 3 identified groups of methods and SWC, CHMM, and DHMM displaying the highest level of similarity.

**Fig. 25.**
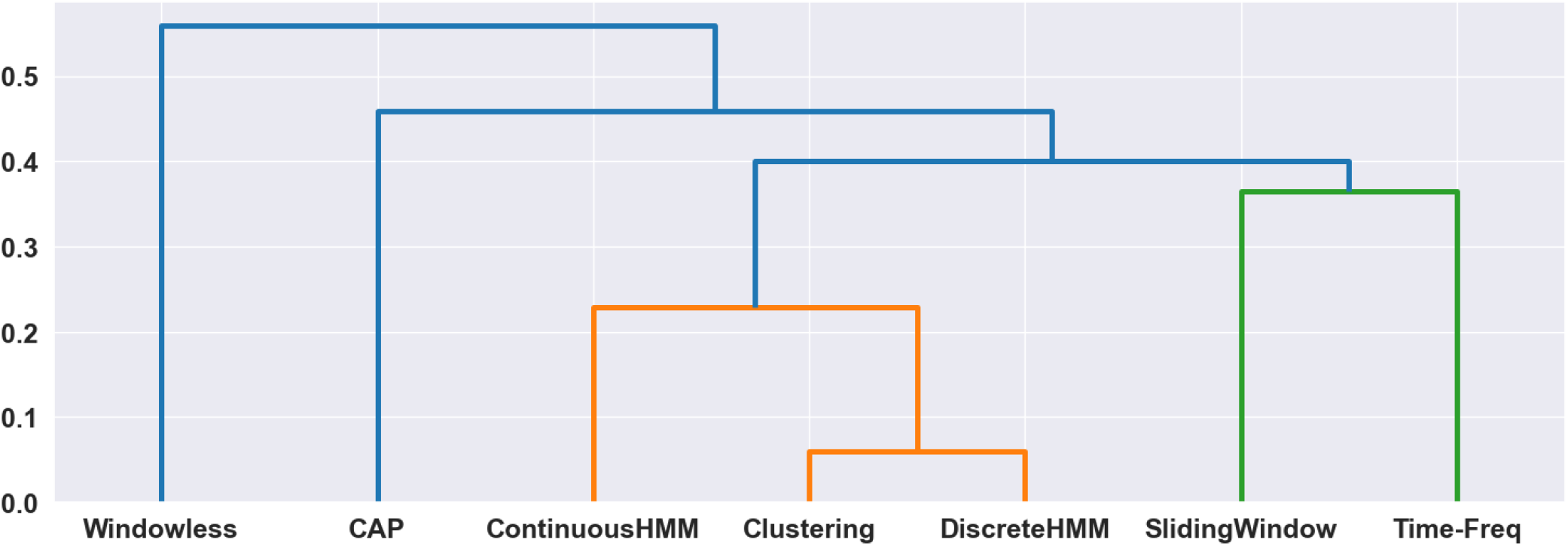
Euclidean distance: Hierarchical clustering of the methods based on the Euclidean distance between their dFC matrices. The dFC matrices were scaled to have unit norm prior to computing Euclidean distances, as the magnitude of dFC matrices are not comparable. The results show a similar pattern of similarity between SWC, CHMM, and DHMM, and between SW and TF similar to the results obtained by Spearman correlation. However, WL exhibits less similarity to other methods, specifically to CAP.

**Fig. 26.**
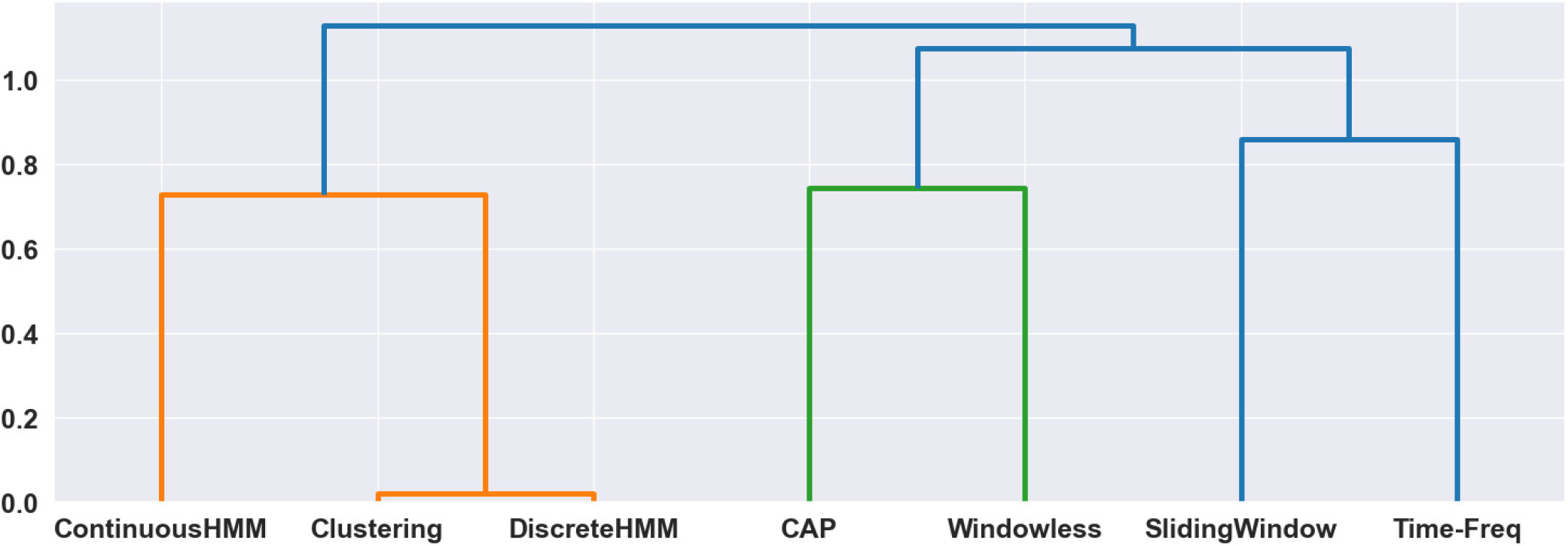
Mutual information: Hierarchical clustering of the methods based on average overall similarity measured by mutual information shows slightly different but overall similar groups of methods compared to the results obtained using Spearman correlation. SWC and DHMM yielded higher mutual information values, while SW shows lower similarity with TF and CHMM shows lower similarity with SWC and DHMM.

**Fig. 27.**
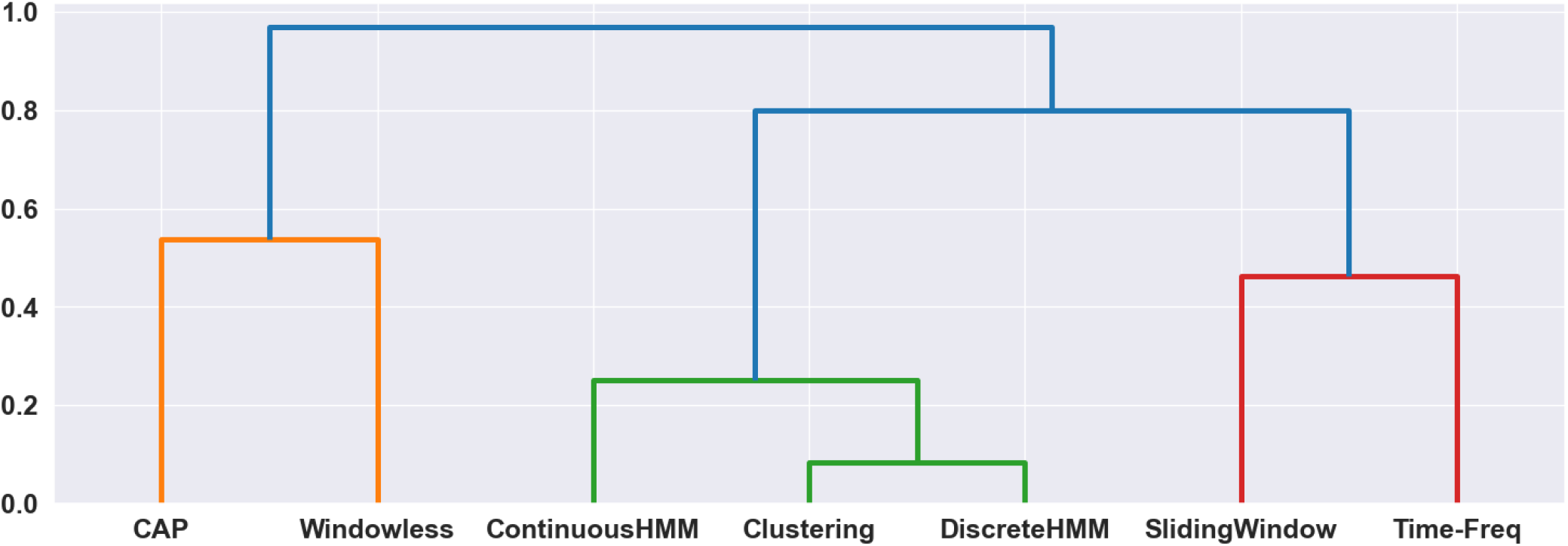
Degree: Hierarchical clustering of the methods based on the degree correlation, as computed by Spearman correlation, of the graphs constructed from their dFC matrices. The correlation values were averaged over time and subjects prior to the clustering analysis. The results yielded the same 3 groups of methods as the other similarity measures.

**Fig. 28.**
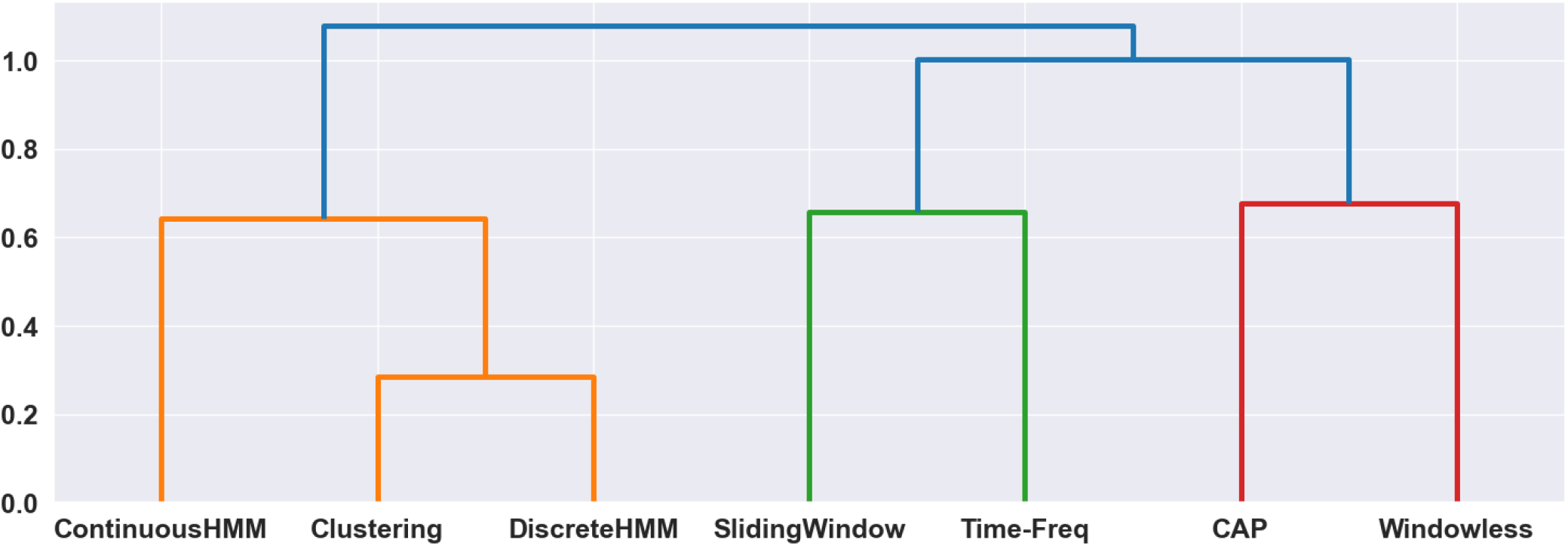
Clustering coefficient: Hierarchical clustering of the methods based on the correlation of the clustering coefficients, as measured by Spearman correlation, of the graphs constructed from their dFC matrices. The correlation values were averaged over time and subjects prior to the clustering analysis. The results, although exhibiting slightly lower similarity values, yielded the same groups of methods as other similarity measures

**Fig. 29.**
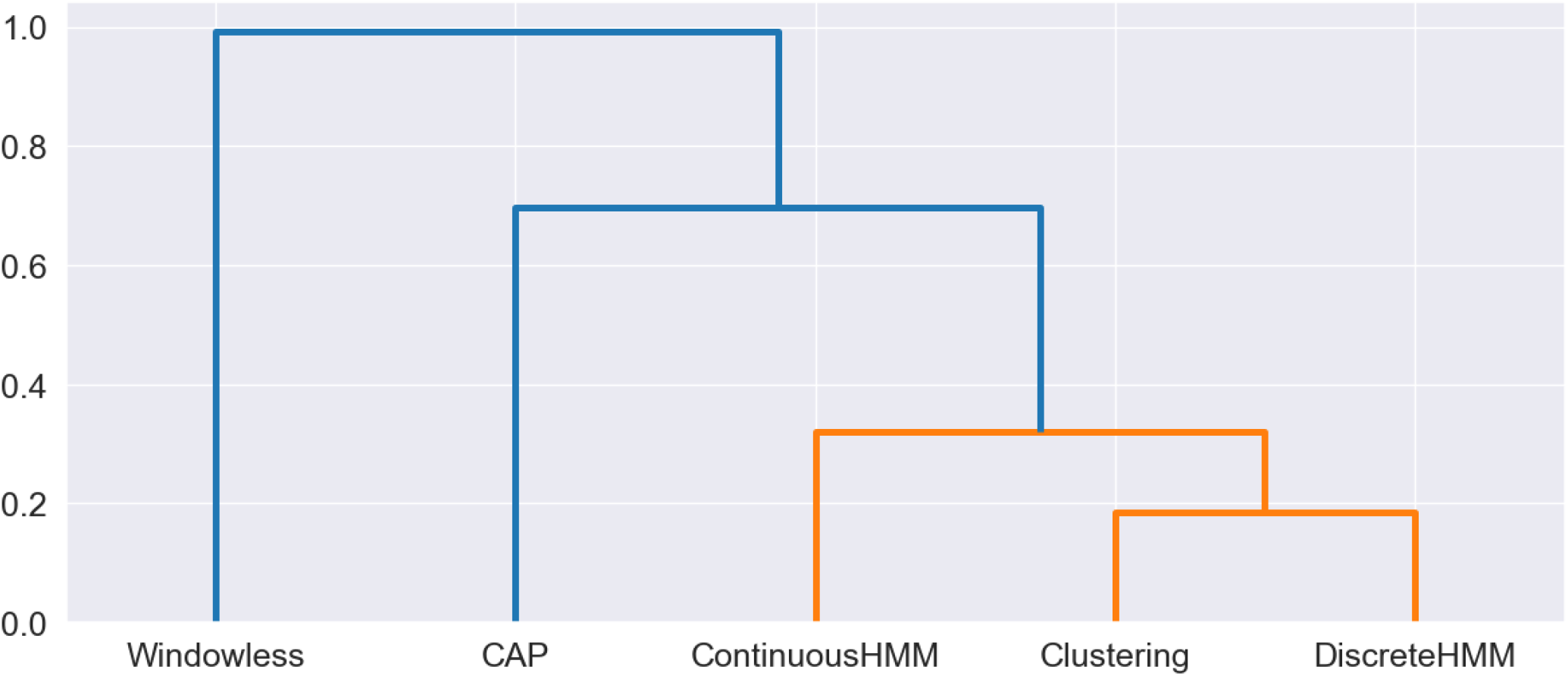
Hierarchical clustering of dFC methods based on inter-subject similarity matrix calculated in terms of the Fractional Occupancy (FO) values derived from state time courses. The inter-subject correlation matrix of each method was measured by computing the Spearman correlation between FO values of different subjects obtained by that method. The results revealed that the inter-subject correlation values obtained by SWC, CHMM, and DHMM were relatively similar to each other, while those obtained by CAP and WL were less similar.

We also compared the variability over method with the variability over subject. After normalizing the dFC matrices by ranking their values, we calculated the variance over method, *var*(*dF C*(*subject*, :, *time, functionalConnection*)), averaging over time and subject, and the variance over subject, *var*(*dF C*(:, *method, time, functionalConnection*)), averaging over time and method. Both calculations resulted in an array of *functionalConnection* values. This yielded an average ratio of *var*_*method*_*/var*_*subj*_ = 0.90, or an average ratio of *SD*_*method*_*/SD*_*subj*_ = 0.94 (see Figure39 in Supplementary material).

## Discussion

### The variability observed in dFC assessment encourages multi-analysis studies

In the present study, our main aim was to assess the analytical flexibility of dFC assessment methods. One of the primary challenges of comparing dFC assessment methods is the lack of a biological reference to determine the significance of the observed variability over methods. While no clear biological reference exists for resting-state fMRI data, we compared the assessed variability over methods with connection variation over time as well as inter-subject variability. The results (Figure4a and Figure39 in Supplementary material) suggest that the variance over methods is on average 0.95 of the variance over time and 0.90 of the variance over subjects. These findings indicate that changing the assessment method may lead to a dFC value which is on average as different as a dFC value obtained at another time point or from another subject using the same method. Moreover, although these biological scales may not be appropriate for every situation, they still can serve as a valid point of reference. Specifically, in the case of comparison with temporal variability, the variation of dFC over time is the main biological phenomenon that the dFC methods aim to measure. These results indicate the variation due to the choice of methodology has an impact as large as the scale of the underlying biological phenomenon of interest.

For further investigation of the analytical flexibility, we conducted a comprehensive comparison of the dFC estimates obtained by different methods using the following approaches:1)subject-wise comparison of dFC matrices in terms of overall similarity (Figure3b); 2) time point-wise comparison of FC matrices in terms of their spatial similarity (Figure3c);3) functional connection-wise comparison of time courses in terms of their temporal similarity (Figure3d); 4) comparison in terms of inter-subject correlation similarity (Figure3e).

Our similarity analysis identified three groups of methods in terms of the similarity of the obtained dFC patterns. The examined methods, while exhibiting fair intra-group similarities, exhibited considerable inter-group variability.

The identified intra- and inter-group similarity profiles can be largely attributed to the underlying assumptions of the examined methods. For example, both WL and CAP disregard the temporal ordering of the BOLD data, allow for instantaneous reconfiguration of FC states, and do not impose a locality assumption. These assumptions enables them to capture more rapid reconfigurations in FC patterns. Conversely, SWC, CHMM, and DHMM, take into account the temporal ordering of the data, and assume locality of neighboring time points, resulting in smoother changes over time (see Figure13 in Supplementary material). These assumptions allow these methods to accurately quantify the temporal dependencies between FC patterns at different time points and the slower dynamics of FC. Lastly, SW and TF, do not assume the presence of re-occurring group-level FC patterns, which allows them to capture the individual subject variability to a greater extent. See Table1 for a summary of the assumptions of each method. These assumptions are rarely invoked in the literature applying dFC technique in a specific study.

The patterns of similarity observed in this study do not imply the superiority of any dFC assessment method or group of methods. Instead, they reveal three groups of methods, each characterized by their specific assumptions, advantages, and disadvantages. These results show that a single analysis or group of analyses may not capture the richness of dFC, and that the use of multiple methods from different groups may reveal different features of dFC. In this regard, developing specialized tools to facilitate the conjoint use of multiple dFC methods, such as the one developed in this study, will aid to fully characterize dFC. For instance, by using one method from the SWC, CHMM, and DHMM group, one from the WL and CAP group, and one from the SW and TF group one can capture rapid FC patterns, as well as temporal dependencies of FC patterns, and subject-specific information.

This approach is particularly important when classifying subjects based on features derived from dFC, as shown in our inter-subject similarity results (Figure3e). We found that using methods from two different groups, e.g., CAP and SWC, can lead to different inter-subject similarity patterns, consequently leading to different clusters of subjects. Hence, we recommend that clustering is performed using multiple methods from different groups. This can help uncover meaningful subgroups or relationships that may not be apparent when using a single method, and ensure that the identified clusters are not solely dependent on the choice of a specific method.

### Understanding temporal complexity in dynamic functional connectivity assessment: what lies behind the variations in dynamic functional connectivity?

Evalu-ating the overall similarity among the outcomes of various methods offers a perspective on the inter-method similarity relationships. However, it remains unclear to what extent these similarities can be attributed to the spatial and temporal aspects of dFC. Based on our results, in most cases, while the spatial comparison of dFC patterns obtained by different methods (Figure3c) exhibited strong similarity between methods, the comparison in terms of temporal dynamics exhibited moderate similarity (Figure3d). Even the methods with the highest spatial similarity, SWC and DHMM, showed low similarity in the time domain. This suggests that capturing the temporal variations of FC may be more challenging than capturing the spatial patterns of dFC. Figure31 in Supplementary material shows that although methods having higher spatial similarity are more likely to have higher temporal similarity, the level of temporal similarity is smaller than the level of spatial similarity in most cases. CHMM, as a particular example, exhibited a high spatial similarity with SWC and DHMM, but a low temporal similarity with these methods (also supported by Figure30 in Supplementary material showing the similarity of inter-time correlations).

**Fig. 30.**
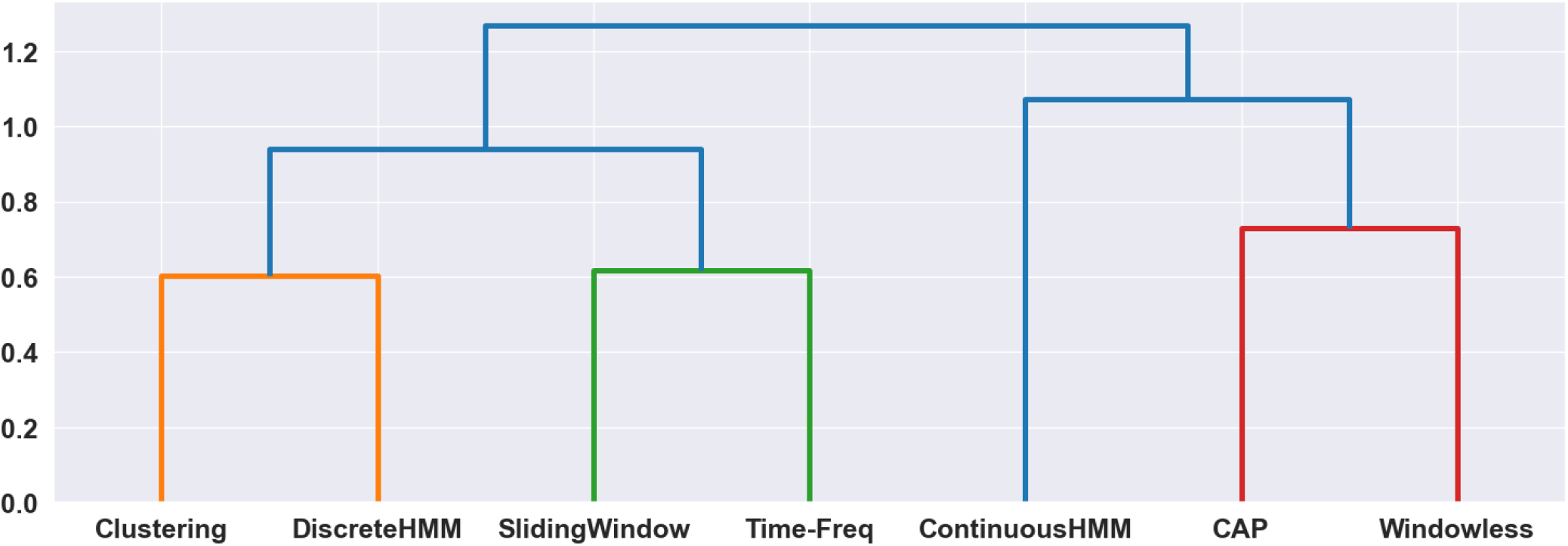
Hierarchical clustering of the methods based on the similarity of their inter-time point correlation matrices, as computed by Spearman correlation. The results indicate a considerable degree of variability across methods, with correlation values generally falling in the weak range. For example, the closest methods based on the overall similarity, such as SWC and DHMM, show weaker correlation compared to when other similarity metrics were used, and CHMM exhibits weak similarity to other methods. However, the overall intra and inter-group similarities, except for CHMM, were similar to those obtained by the overall similarity.

**Fig. 31.**
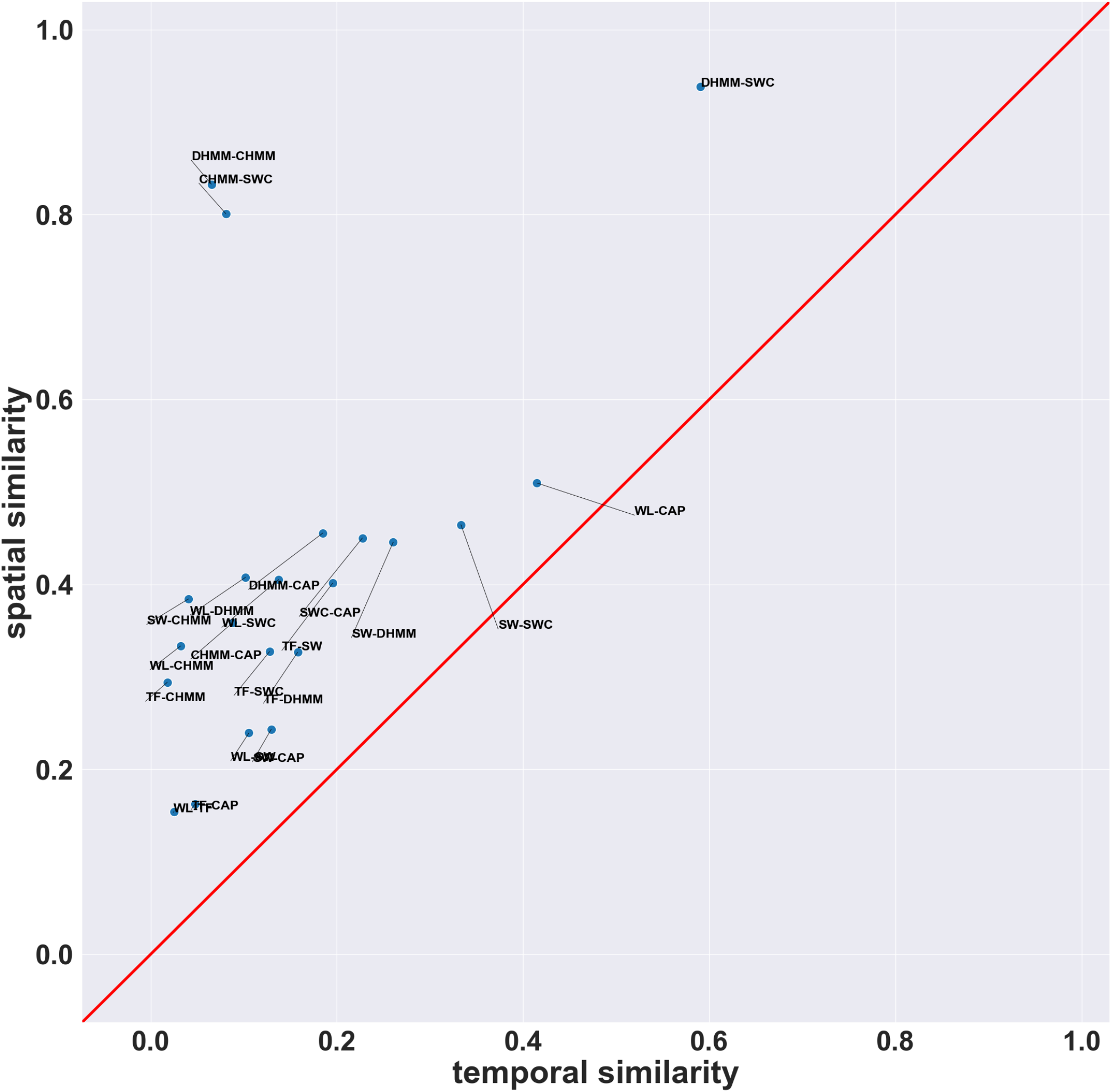
Scatter plot comparing the spatial and temporal similarity of the examined dFC assessment methods. The majority of the methods show a linear relationship between their spatial and temporal similarity, with higher temporal similarity corresponding to higher spatial similarity. However, the temporal similarity generally tended to be weaker compared to spatial similarity. Notably, DHMM-SWC exhibited the highest levels of both spatial and temporal similarity, while DHMM-CHMM and CHMM-SWC had high spatial similarity but low temporal similarity. Additionally, WL-CAP, despite having an average spatial similarity, had a higher temporal similarity compared to most of the other pairs.

Our supplementary material results suggest that in several cases, pairs of methods did not exhibit a significantly greater similarity compared to the similarity obtained when the dFC matrices were randomly shuffled with respect to time (Figure32; note that the fractional occupancy of the states was preserved for the shuffled dFC matrices). Pairs of methods exhibiting strong similarity between their time-shuffled dFC matrices indicate that the average similarity between all pairs of their spatial patterns, or FC matrices, even at different time points, is high. For instance, CHMM-SWC exhibited a relatively high overall similarity, however, their similarity did not change significantly when the dFC matrices were time-shuffled. This implies that the dFC matrices of CHMM and SWC did not exhibit more consistency in terms of their patterns of variation over time than the time-shuffled dFC matrices and that the observed strong similarity between them was mostly a result of the strong similarity between their spatial patterns. On the other hand, WL-CAP, despite having a low overall similarity, showed significantly greater similarity when compared to the shuffled-time similarity distribution. This implies that their temporal variations exhibited more consistent patterns than those of the time-shuffled dFC matrices, indicating that their low overall similarity was caused by low similarity of their spatial patterns. These observations highlight that the variability in the dFC results may spread differently across the temporal and spatial dimensions, and strong similarity in one dimension does not necessarily imply similarity for the other dimension. It also suggests that comparing the overall similarity of dFC results, or comparing only their spatial patterns may not provide a comprehensive understanding of the relation between methods.

**Fig. 32.**
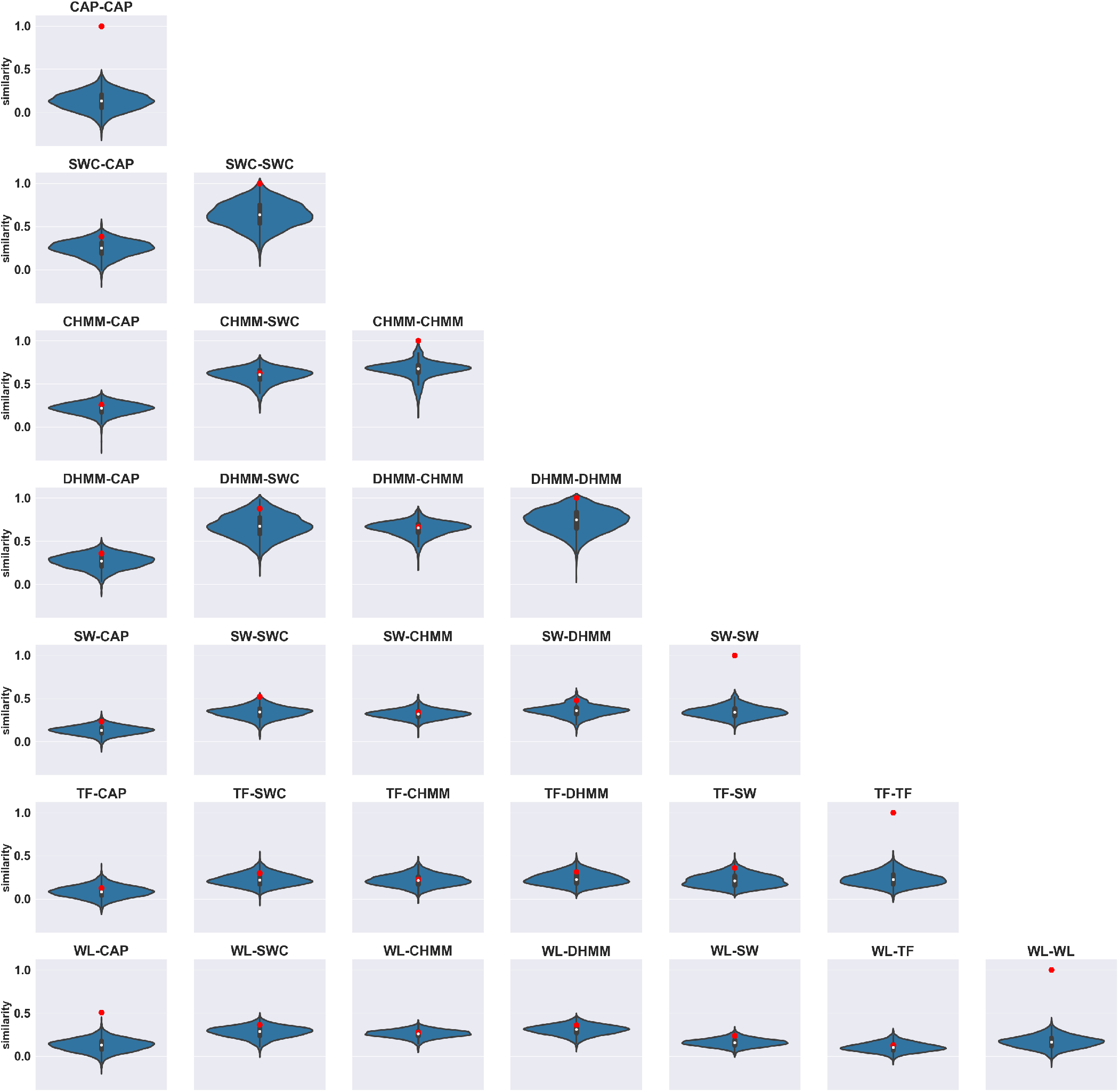
The violin plots display the distribution of overall similarity between dFC matrices for each method pair after 10 different random shufflings of the time points for each subject (totaling 3,950 samples). The red dots indicate the average of actual similarities over subjects. For certain pairs, such as WL-CAP, DHMM-SWC, and SW-SWC, the average values of actual similarity were significantly higher than the average similarity of shuffled dFC matrices, implying that the temporal alignment or synchrony of dFC matrices was significant. This observation for pairs such as WL-CAP also suggests that their low overall similarity is likely due to dissimilar spatial patterns rather than dissimilar temporal dynamics. Conversely, for other pairs like DHMM-CHMM, the average values of similarity were not significantly greater, suggesting that their overall similarity was mainly attributed to the similarity of their spatial patterns, or due to their smooth transitions, and that their temporal similarity did not significantly contribute to the overall similarity. The distributions were estimated using the kernel density estimation (KDE) algorithm.

**Fig. 33.**
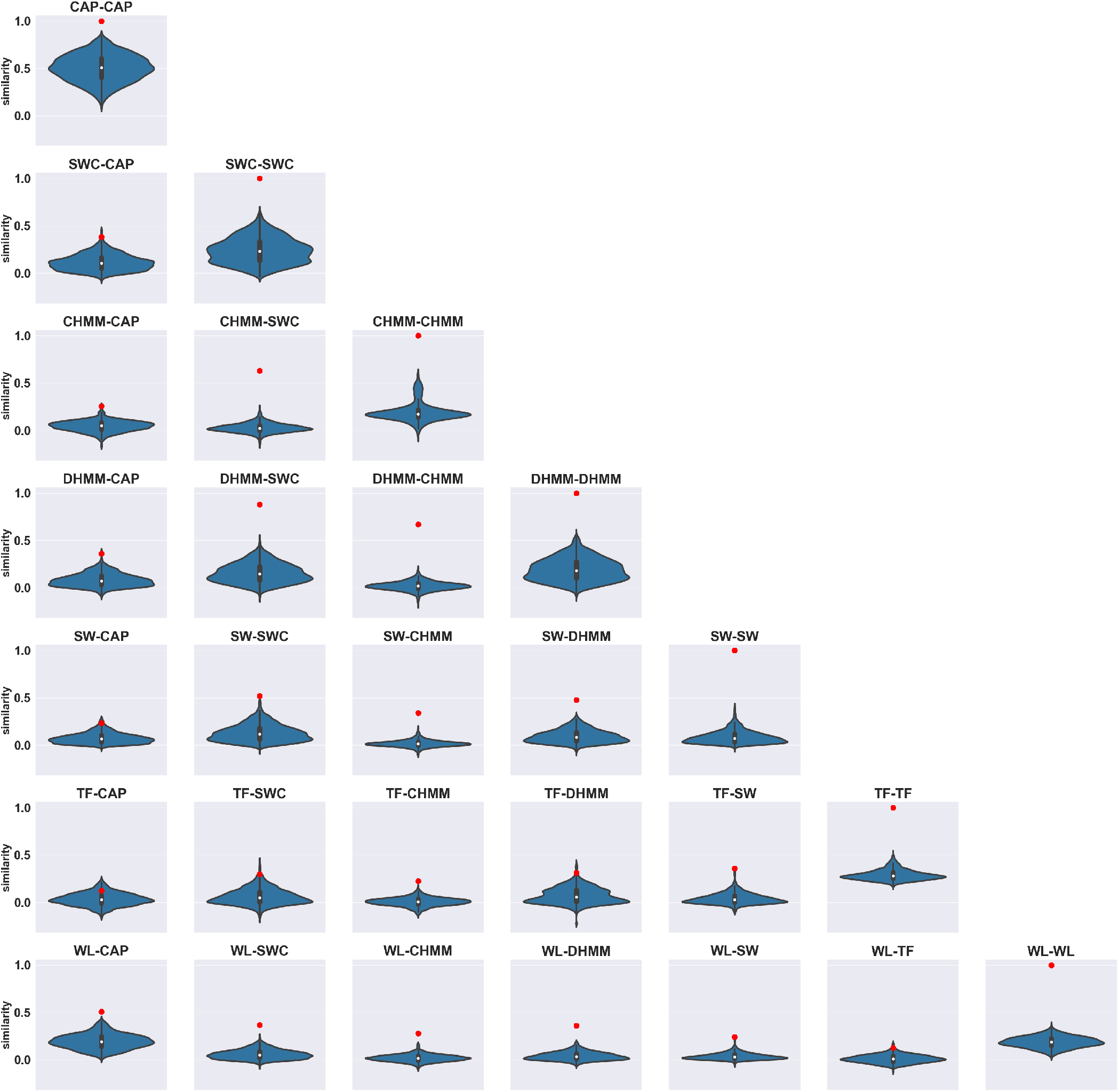
The violin plots display the distribution of overall similarity between dFC matrices for each method pair after 10 different shuffling of regions, and hence functional connections, for each subject (totaling 3,950 samples). The red dots indicate the average of actual similarities over subjects. For all pairs the average value of similarity was greater than the average similarity of shuffled dFC matrices, implying that their spatial patterns, or FC matrices, at each time point were more similar than the average of randomly shuffled dFC matrices. For certain pairs, such as CHMM-SWC, DHMM-SWC, and DHMM-CHMM, the observed similarity values greatly exceeded those of the shuffled dFC matrices, indicating that their spatial patterns are significantly more similar than those generated randomly. Conversely, for pairs such as TF-CAP and WL-TF, the actual similarity values were comparable to the randomized similarity values. The distributions were estimated using the kernel density estimation (KDE) algorithm.

**Fig. 34.**
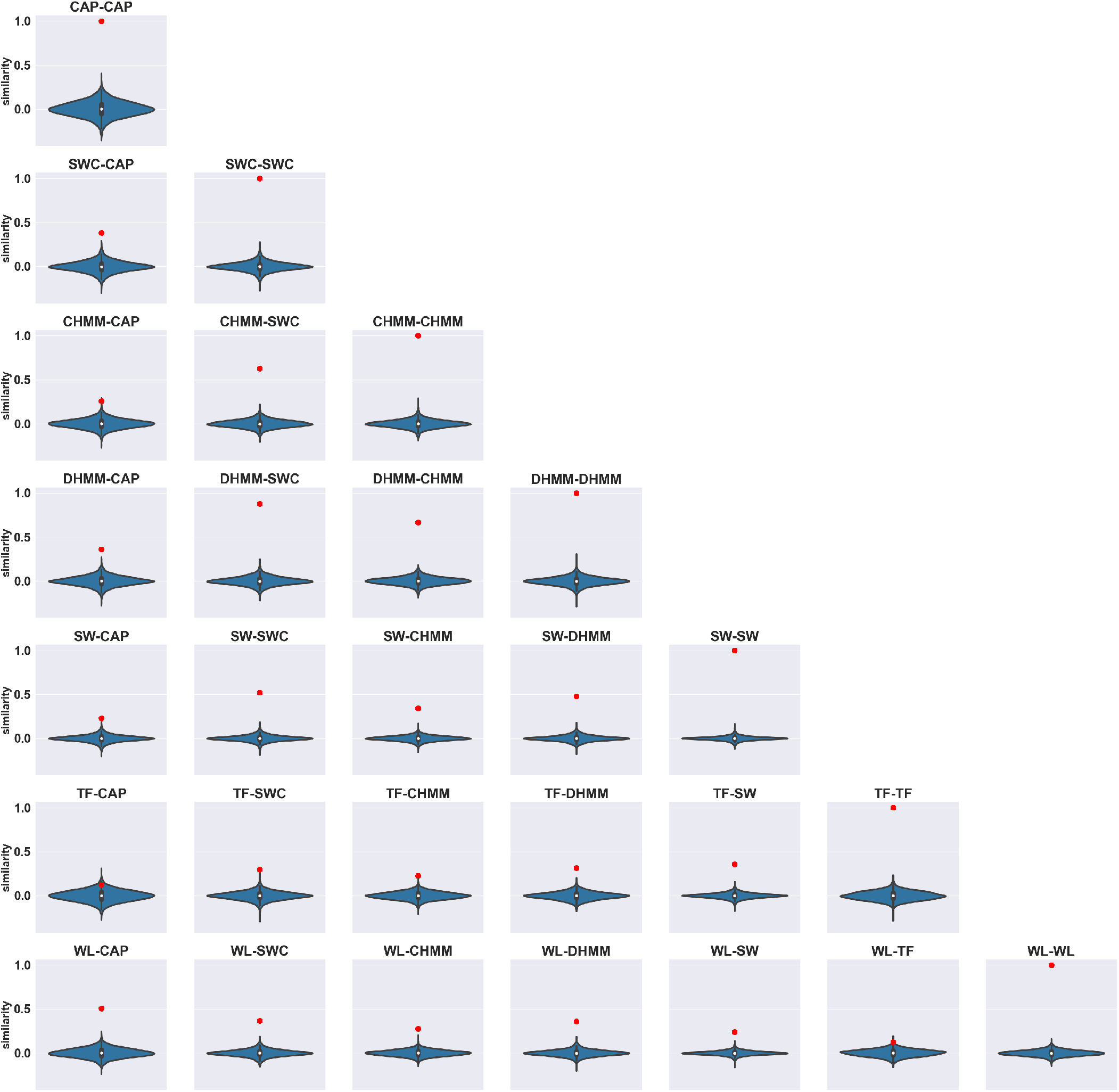
The violin plots display the distribution of overall similarity between dFC matrices for each method pair after 10 different random shufflings of both time points and functional connections for each subject (totaling 3,950 samples). When both time points and functional connections were shuffled, both the spatial and temporal patterns of dFC matrices were removed. The red dots indicate the average of actual similarities over subjects. The results suggest that for all pairs of methods the actual similarity was greater than the average similarity of shuffled dFC matrices, implying that dFC matrices of all pairs were more similar than two dFC matrices with randomized temporal and spatial patterns. However, for certain method pairs such as TF-CAP and WL-TF, the actual similarity values were comparable to the average similarity of random dFC matrices. The distributions were estimated using the kernel density estimation (KDE) algorithm.

**Fig. 35.**
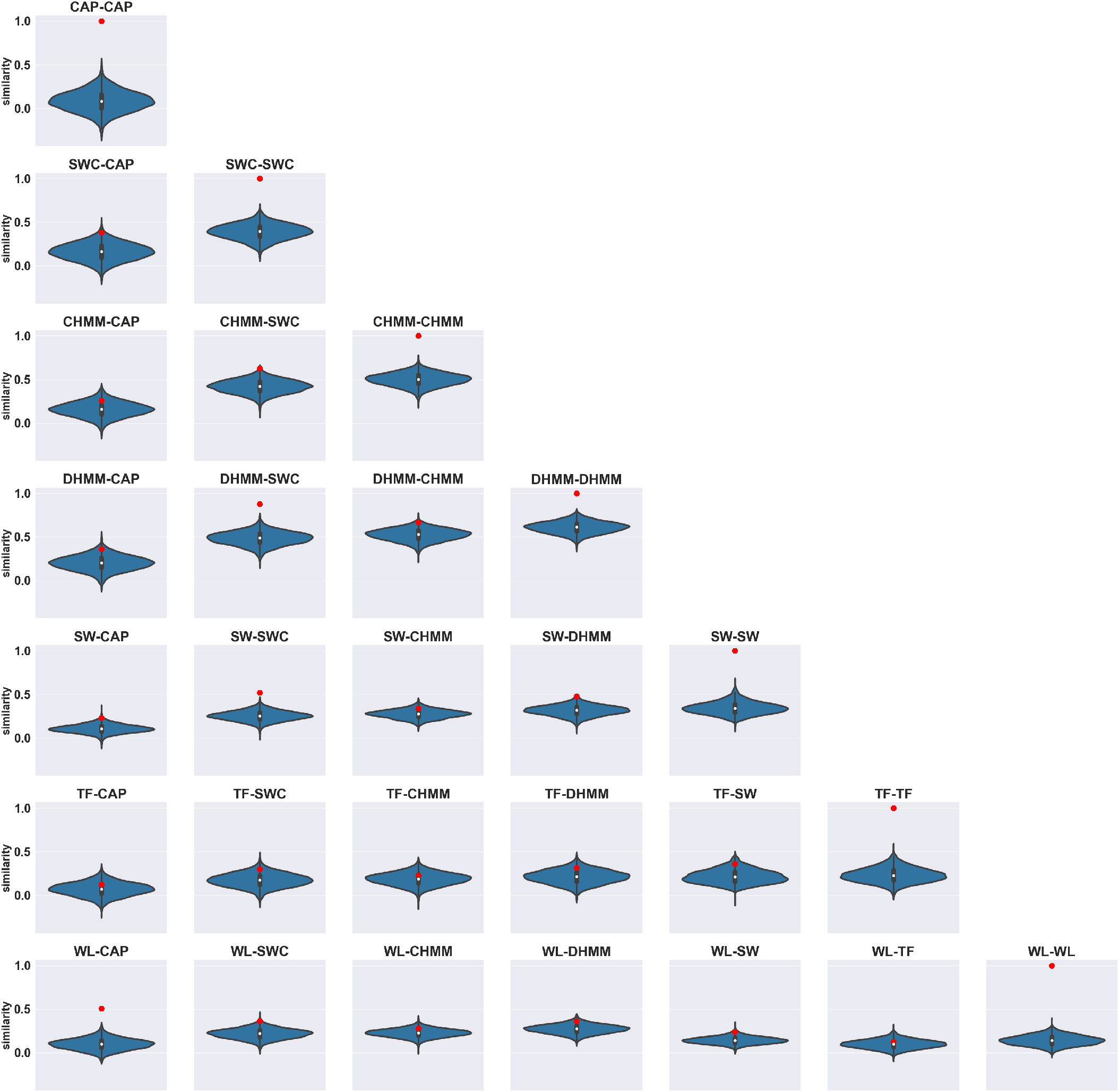
The violin plots depict the distribution of overall similarity between randomly generated dFC matrices. The randomly generated dFC matrices of each method were obtained by generating a random state time course and using the obtained spatial patterns of the method. For each subject, 10 random dFC matrices were generated by using the actual FC matrices, or spatial patterns, of each method, resulting in 3,950 samples for each method pair. For SW and TF, all spatial patterns occurring in the subject’s dFC matrices were used (38 patterns). The red dots show the average of actual similarities over subjects. For certain pairs such as WL-CAP and DHMM-SWC, the average value of the actual similarity was significantly higher than the average similarity of randomly generated dFC matrices, indicating that the alignment of their dFC matrices was significantly stronger than two randomly generated dFC matrices with the same spatial patterns but random state time courses. Conversely, for pairs such as DHMM-CHMM, the same comparison suggests that any random sequence of the same spatial patterns may have a similar level of overall similarity as their actual dFC matrices. This implies that a large portion of their overall similarity stems from the similarity between their spatial patterns rather than the smoothness of transitions or temporal synchrony. The distributions were estimated using the kernel density estimation (KDE) algorithm.

**Fig. 36.**
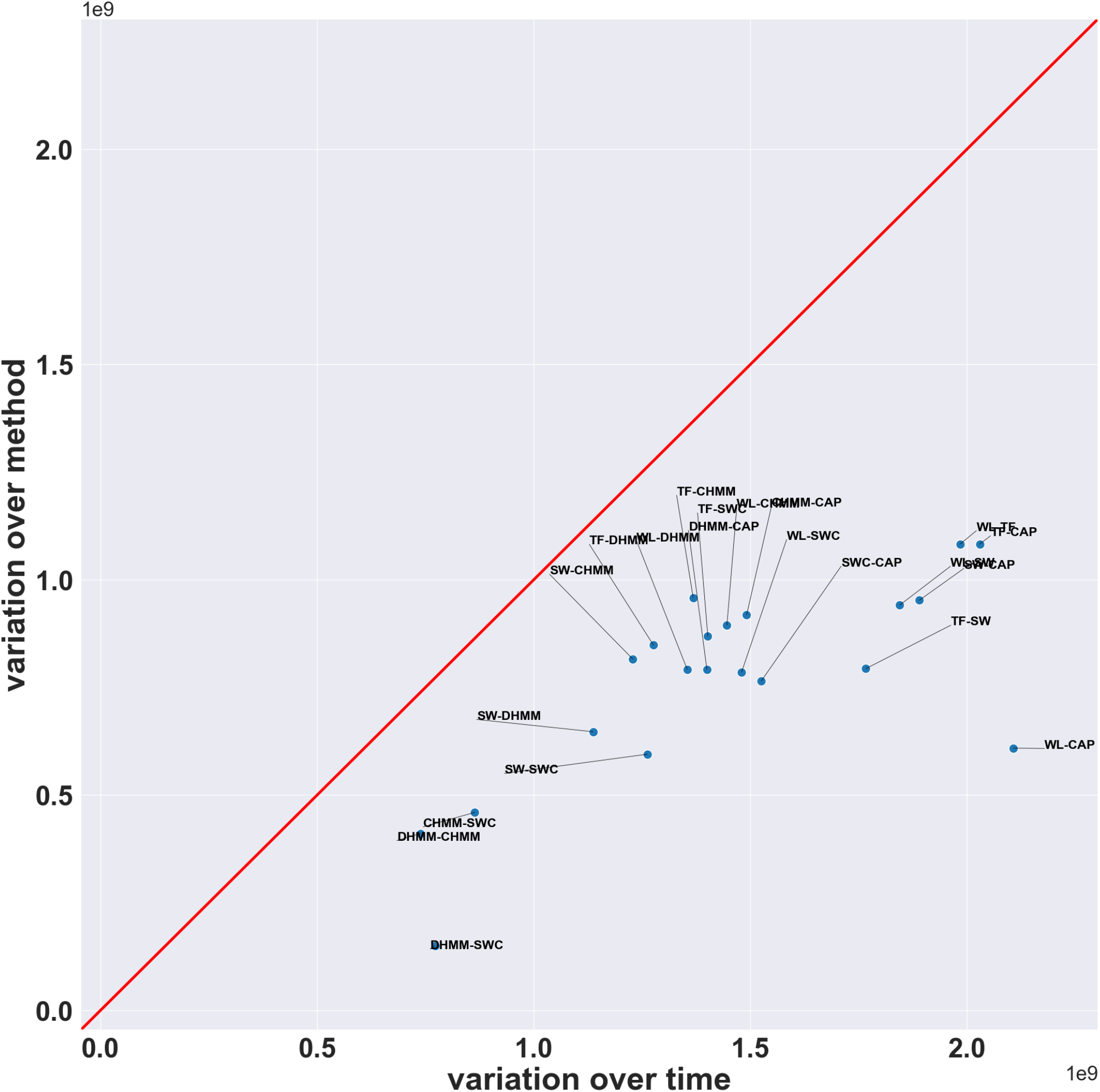
Scatter plot comparing the variance of dFC matrices over the time axis and the variance of dFC matrices over the method for each pair of methods. To account for the different value distributions of each method, the dFC matrices for each subject, *dF C*(*subject*_*n*_, *method*_*i*_, *time, ROI, ROI*), were normalized by ranking the values, similar to the approach used when calculating Spearman correlation. For each pair, the dFC matrices were concatenated, *dF C*(*subject*, [*method*_*i*_, *method*_*j*_], *time, ROI, ROI*). Their variance over time and method were calculated by computing variance over the time axis and method axis respectively, and averaging over the other axis. The pairs DHMM-SWC, CHMM-SWC, and DHMM-CHMM exhibited both the lowest variance over time and over method, while WL-TF and TF-CAP yielded high variance over both time and method. Notably, WL-CAP, despite having the highest variance over time, yielded an average variance over method.

**Fig. 37.**
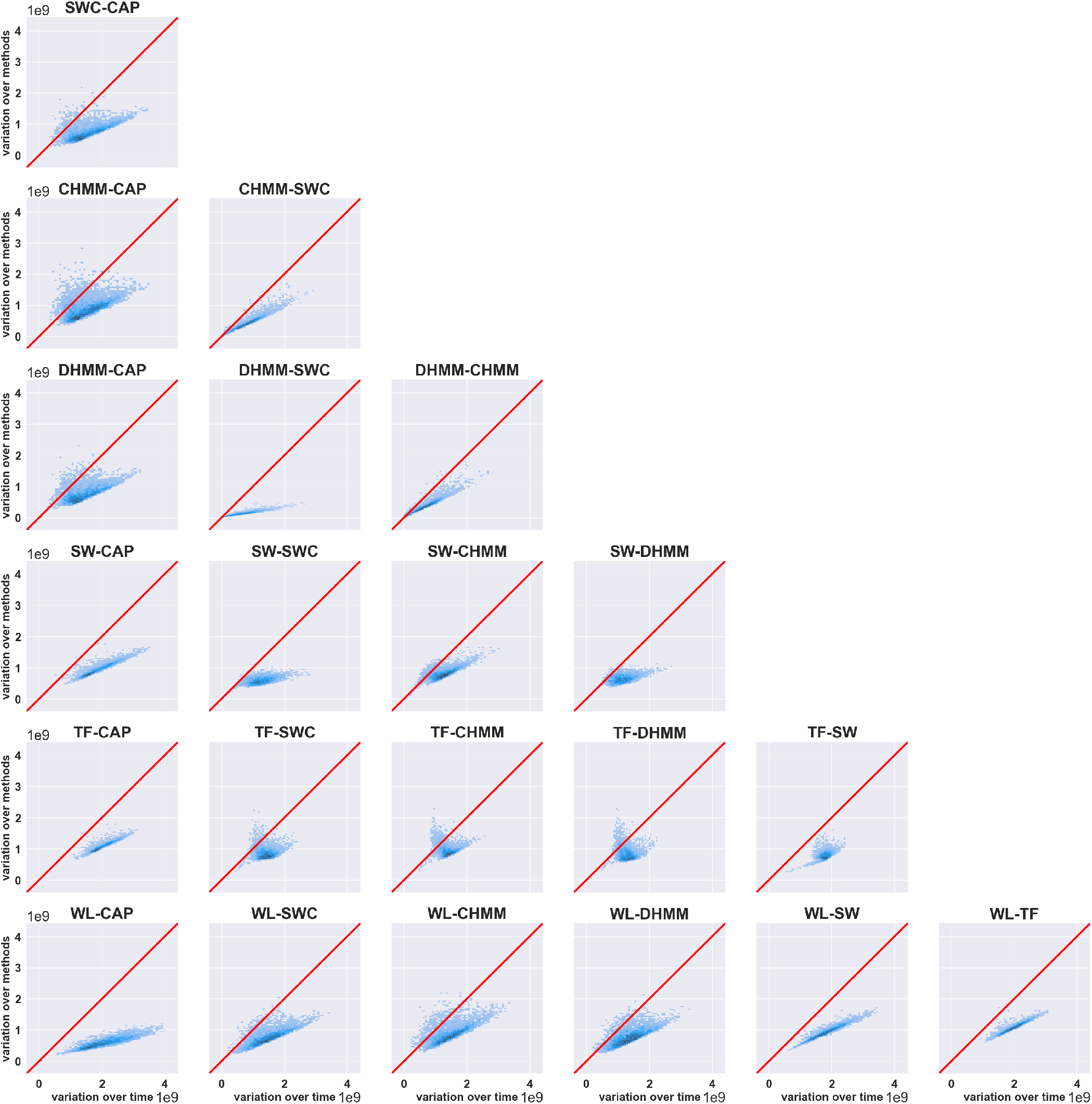
Scatter plots of the variance over method versus the variance over time for each functional connection across all subjects (similar to Figure4a but created for each pair of methods separately). To account for the different value distributions of each method, the dFC matrices for each subject, *dF C*(*subject*_*n*_, *method*_*i*_, *time, ROI, ROI*), were normalized by ranking the values, similar to the approach used when calculating Spearman correlation. For each pair, the dFC matrices were concatenated, *dF C*(*subject*, [*method*_*i*_, *method*_*j*_], *time, ROI, ROI*). Their variance over time and method was calculated by computing the variance over the time axis and method axis respectively, and averaging over the other axis. The scatter plots reveal a wide range of behaviors across method pairs. For some pairs, such as DHMM-SWC and WL-CAP, the ratio between variation over method divided by temporal variation was low, while for pairs such as WL-DHMM and TF-DHMM, the ratio was closer to one.

**Fig. 38.**
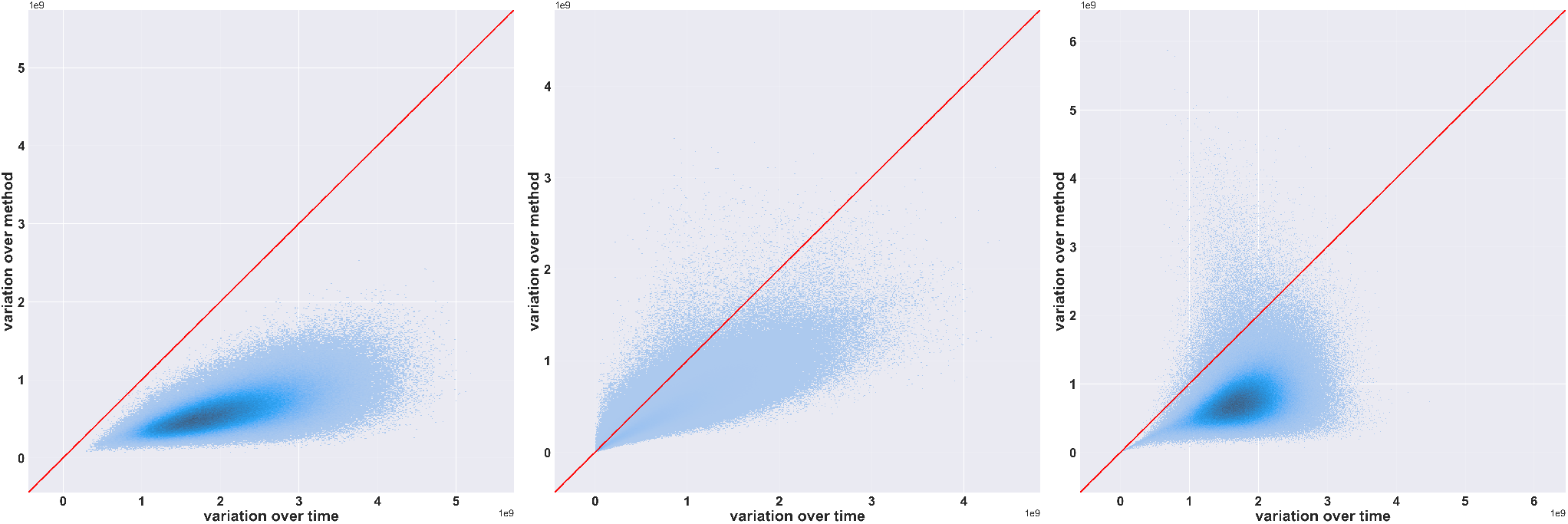
Scatter plots of the variance over method versus time for each functional connection across all subjects similar to Figure4a but created for each group of methods separately: Group 1) CAP and WL; Group 2) SWC, CHMM, and DHMM; Group 3) SW and TF. The ratios were: Group 1: 0.31, Group 2: 0.65, Group 3: 0.46.

**Fig. 39.**
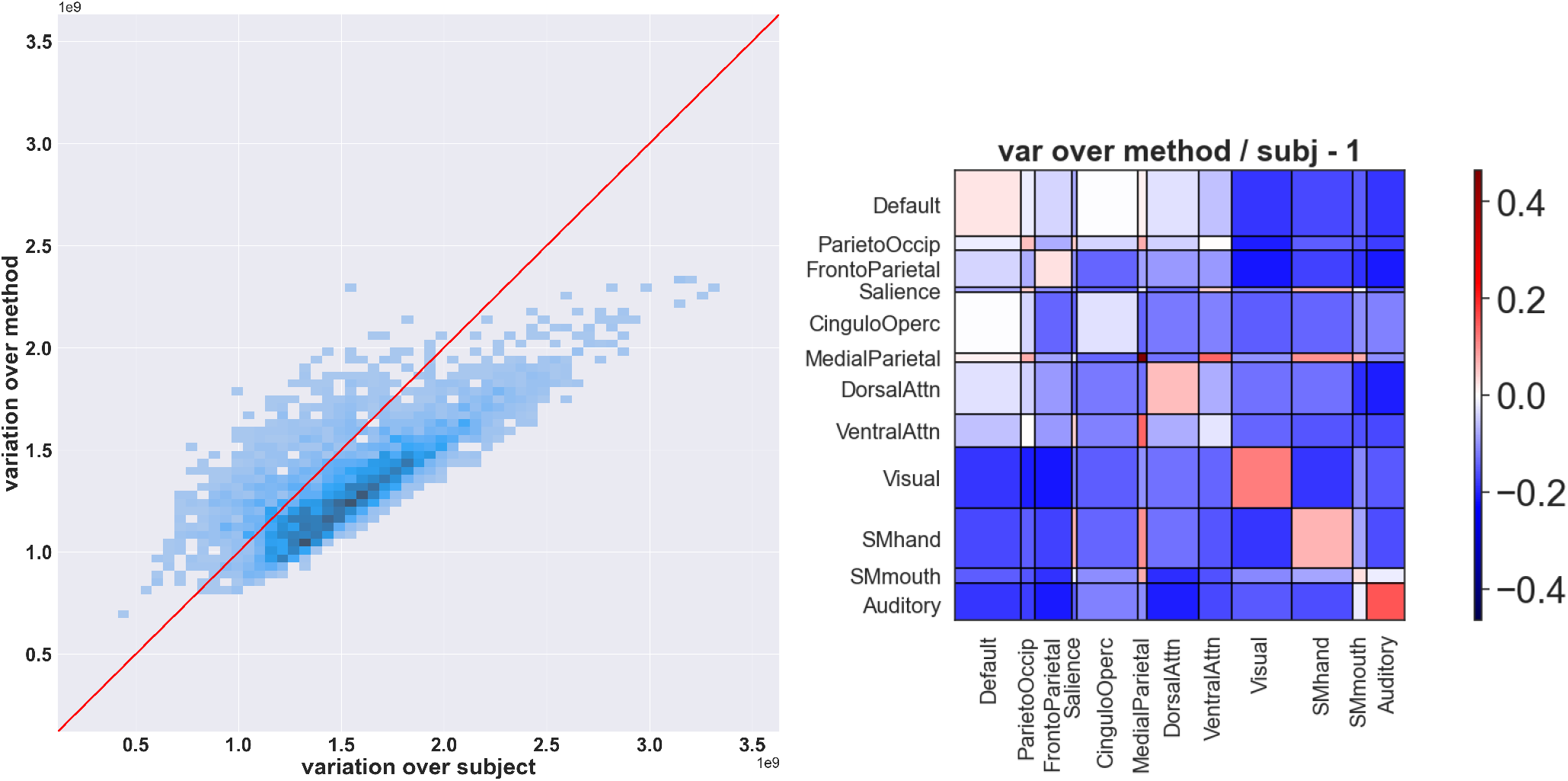
**Left:** Scatter plot demonstrating the comparison of variance over methods and variance over subjects for each functional connection, averaged over time ((*ROI ×* (*ROI −*1)/2) points in total). The results reveal that the variance observed across dFC methods is comparable to the variance observed across subjects, with an average ratio of *var*_*method*_*/var*_*subj*_ = 0.90 (*SD*_*method*_*/SD*_*subj*_ = 0.94). **Right:** Ratios of variance over methods divided by variance over subjects across functional connections and averaged over time. To facilitate comparison, the ratios were subtracted by one (*ratio −* 1), with zero values (white) representing equal variance over methods and subjects. The figure highlights pairs of RSNs with functional connections that exhibited higher variability over methods (red), as well as RSNs that exhibited higher variability over subjects (blue). Prior to computing the ratios, the variance values for functional connections belonging to each RSN pair were averaged to obtain a single variance value. The plot indicates that functional connections between RSNs such as the Default Mode and Parieto-occipital Networks, Default Mode and Cingulo-Opercular Networks, as well as those within the Default Mode, FrontoParietal and Cingulo-Opercular Networks demonstrated relatively equal variance over subjects and methods. Conversely, functional connections within RSNs, such as the Visual and Auditory networks, as well as those between the Ventral Attention and Medial Parietal Networks, exhibited higher variation over methods compared to subjects.

**Fig. 40.**
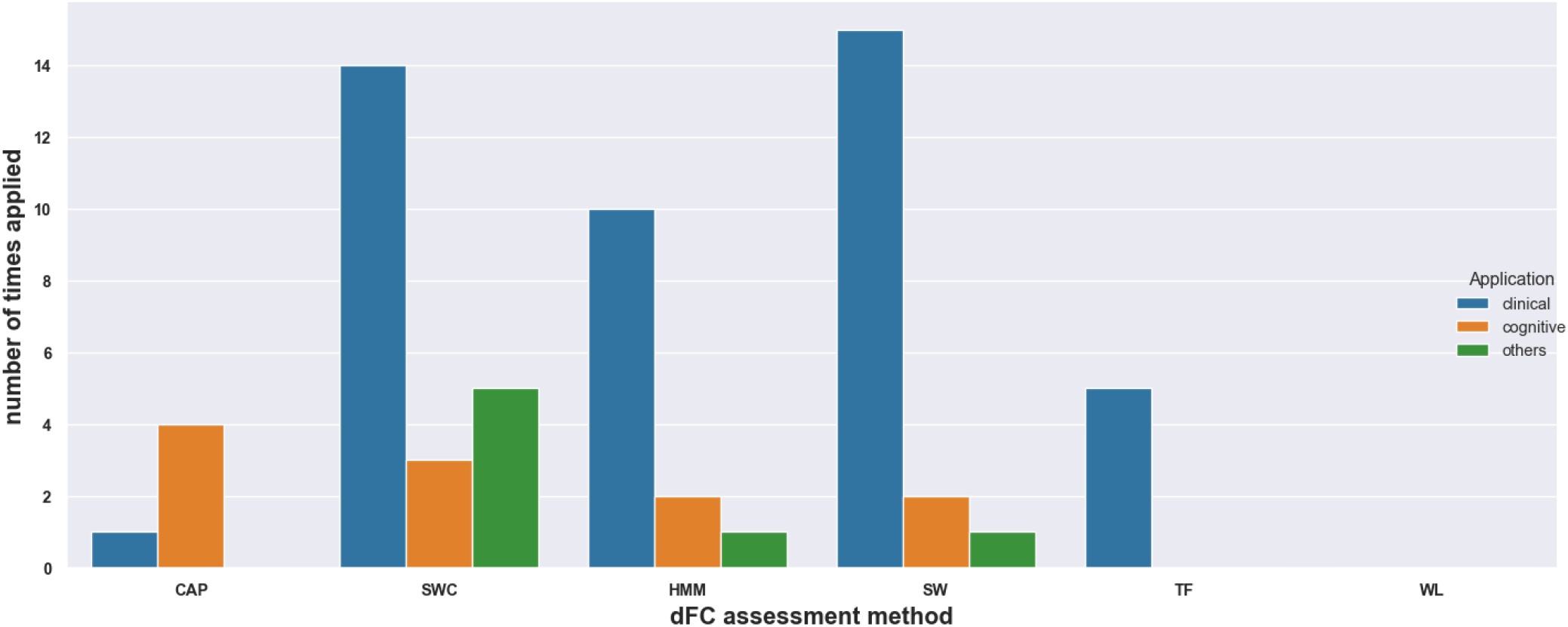
Bar plots showing the number of times each dFC method was used in each application across around 200 papers that were reviewed. The bar plots show that SW and SWC were the most popular methods in papers related to clinical applications and CAP was the most popular method in papers related to cognitive application.

Future research should conduct more in-depth comparisons of dFC assessment methods, focusing on the temporal dynamics of their outputs, to shed light on the reason for the observed variability over methods. This is crucial, as temporal dynamics play a significant role in dFC and have already been a source of controversy regarding their origins. Our results suggest that a part of these temporal variations may originate from the methodology used for assessing them.

Previous studies have demonstrated that physiological factors, such as respiration, cardiac activity, and motion artifacts, also modulate the temporal variations of dFC [47, 48]. These non-neural sources of variability can confound the interpretation of dFC patterns and potentially lead to erroneous conclusions. Future research could develop advanced techniques, including denoising algorithms, physiological signal regression, and data-driven approaches, to effectively isolate the neural-driven part of dFC variations from the physiological sources. By disentangling these sources, researchers could gain a more precise understanding of the genuine neural dynamics underlying dFC, enhancing the reliability of dFC studies and their potential applications in clinical diagnostics and treatment evaluations.

### Study limitations and future directions

Potential future work should review external validation techniques and develop a validation framework to assess the accuracy of dFC assessments in more controlled paradigms, such as predicting cognitive state of the subjects, using simulated data, and assessing the consistency of the result with the neural activation patterns derived from simultaneous electrophysiological recordings. Evaluating the dFC assessment methods using such paradigms will lead to a better understanding of the reasons behind the observed variability between the results yielded by different dFC methods. Furthermore, it would shed light on the varying levels of method variability observed across different RSNs by identifying the factors that contribute to higher variability in certain RSN pairs compared to others. Additionally, this would also provide a more reliable biological reference for quantifying the magnitude of the observed methods’ variability. One of the limitations of the current study is the lack of a clear biological reference. The nature of resting-state data makes it challenging to establish expected patterns or variations in dFC over time or across subjects. This is primarily because subjects are free to engage in any mental activity during the resting state [4, 60]. In contrast, controlled paradigms, such as the ones mentioned earlier, involve specific instructions or tasks, resulting in more deterministic dFC patterns. For instance, in task-based paradigms, subjects perform specific tasks, leading to more comparable dFC patterns and enhanced interpretability of inter-subject variability. Similarly, simultaneous fMRI and electrophysiological recordings can provide a more reliable biological reference through the integration of electrophysiological activity. Therefore, future work should prioritize the use of such paradigms to facilitate the utilization of more interpretable biological references.

It is worth noting that in our study, we adopted the hyperparameter values recommended by the original paper or consensus among the community for each method, which may contribute to the observed variability in our results. Different hyperparameter choices could lead to different similarity patterns among methods and varying levels of variability. Therefore, future work should also investigate the sensitivity of dFC assessment results to the choice of hyperparameters and their effect on the observed variability.

Assessing the robustness of neuroscience results with respect to dFC analytical flexibility should be strongly recommended. We propose the utilization of multi-analysis approaches, preferably using methods that show less similar results, for assessing dFC and reporting a more comprehensive view of the data, made easier with the python toolbox released with this article. While the use of several methods may cause interpretation and statistical challenges, assessing the variability of the results is important. As this may not always be feasible, it is also reasonable to select a single dFC method based on the specific research objectives, method strengths and limitations, compatibility with the study design and data characteristics.

We have developed a Python toolbox, termed `multi_analysis_dfc`, that simplifies the implementation of multi-analysis dFC assessment. The toolbox enables users to apply various dFC assessment methods to their data and experiment with different hyperparameters, allowing for a comprehensive analysis of dFC variability over methods and hyperparameters. Users can compare the results of different methods within a single package, making it easy to explore and evaluate dFC assessment techniques. This toolbox is unique in providing an open-source implementation of commonly used dFC methods and should be a valuable resource for researchers seeking to explore and compare different dFC assessment techniques. The toolbox is available on GitHub and is designed to be a collaborative package where other methods implementation can be integrated.

## Conclusions

In conclusion, the present study aimed to evaluate the analytical flexibility of dFC measurement methods and establish how analytical flexibility compares to biological variability. The comparison results show a wide range of similarity patterns across methods. The variability found in the results highlights the importance of carefully motivating and validating the choice of a specific dFC assessment method. The use of multiple methods would mitigate the issues arising from analytical flexibility and better characterize the richness of dFC measurements. Future work should aim to develop a validation framework and evaluate the accuracy of dFC assessments through external validation techniques.

## Abbreviations

AD: Alzheimer’s Disease
ADHD: Attention-Deficit/Hyperactivity Disorder
ASD: Autism Spectrum Disorders
BOLD: Blood-Oxygen-Level-Dependent
CAP: Co-Activation Patterns
CHMM: Continuous Hidden Markov Model
dFC: dynamic Functional Connectivity
DHMM: Discrete Hidden Markov Model
DOC: Disorders Of Consciousness
FC: Functional Connectivity
fMRI: functional Magnetic Resonance Imaging
FO: Fractional Occupancy
HCP: Human Connectome Project
HMM: Hidden Markov Model
KDE: Kernel Density Estimation
LR: Left-to-Right
MCI: Mild Cognitive Impairment
PCA: Principal Component Analysis
PD: Parkinson’s Disease
PS: Phase Synchrony
PTSD: Post-Traumatic Stress Disorder
RL: Right-to-Left
ROI: Region Of Interest
RSN: Resting State Network
SD: Standard Deviation
SW: Sliding Window
SWC: Sliding Window + Clustering
SZ: SchiZophrenia
TF: Time-Frequency
t-SNE: t-distributed Stochastic Neighbor Embedding
WL: Window-Less
WTC: Wavelet Transform Coherence

## Supplementary material

In this document, we present supplementary data that offers additional insights into the implemented methods, as well as additional comparisons and analyses to evaluate their similarity and the variability of dFC results. These supplementary findings provide valuable additional information to complement the main study and further enhance our understanding of the assessed methods and their outcomes.

## Additional information regarding individual methods

This section provides supplementary information and descriptions for each individual method used in the study. It offers further insights into the specific characteristics, assumptions, and dFC results.

### state FC matrices

The 12 state FC matrices of each implemented methods. Each state FC matrix is *ROI*× *ROI* and ROIs are divided into 12 RSNs. The state FC matrices were each normalized to (− 1, 1) for better visualization. For each state, the mean brain activity across ROIs is also shown. Note that the state FC matrices in these figures are denoted using the term ‘FCS.’

### Average dFC (static FC)

### Temporal variance of dFC

### Inter-state spatial similarity in state-based methods

### Similarity of adjacent time points or smoothness

### Frequency of inter-state transitions in state-based methods

### Dwelling times of state-based methods

### Computation time: fitting states

### Computation time: estimating dFC

## dFC Values Visualization

This section provides visualizations of the dFC values obtained from different methods, offering insights into their distributions across subjects and methods.

### Distributions of dFC values

### Embedded Presentation of subjects dFC

## More in-depth exploration of the similarity between dFC assessment methods

This section contains supplementary data and analyses that offer additional insights and detailed information about the similarity assessments conducted in the main study.

### Distribution of the similarity of dFC assessment results obtained by Spearman correlation over subjects

### Hierarchical Clustering Structure across subjects

### Results of two-way ANOVA test on the effect of session (day and direction) on the measured similarities

The two-way ANOVA test was performed separately for each pair of methods to investigate the effect of day and direction of the scan on their similarity.

### Similarity across functional connections

### Overall similarity using other metrics

### Similarity assessment based on graph properties

To evaluate the similarity of dFC matrices obtained by various methods in terms of graph-based properties, we constructed a dynamic graph of the brain using the FC matrix of each time point. The ROIs were represented as nodes and the functional connections between them as edges, resulting in a sequence of graphs over time. We then computed a set of graph properties for each graph and compared them across methods to assess the similarity of the methods in terms of these properties.

### Inter-subject similarity by Fractional Occupancy (FO) feature

Fractional Occupancy (FO) is a measure of the fraction of time spent at each state by dFC matrices obtained by state-based methods. This feature has become popular as a dFC-based feature for classifying subjects. Here, we used FO values obtained using the state time courses of different state-based methods to obtain the inter-subject correlations. Note that since this feature is based on state time courses, it is not applicable to state-free methods.

### Correlation of inter-time point similarity patterns

To assess the relationship between FC patterns at different time points in a dFC matrix, we constructed an inter-time point correlation matrix. Each entry (*i, j*) in the matrix represents the correlation between the dFC values at time point *i* and time point *j*. The diagonal entries represent the correlation of each FC matrix at time point *t*, or *dFC*(*t*), with itself, while the off-diagonal entries show the correlation between the FC matrices at different time points, e.g. *dFC*(*t*_*i*_) and *dFC*(*t*_*j*_).

## Spatial vs. Temporal

This section contains supplementary data and visualizations related to the spatial and temporal similarity assessment conducted in the main study. It includes comparison of spatial and temporal similarity values and additional analyses with randomization tests to compare the actual similarity values, obtained from the original dFC matrices, with those obtained using dFC matrices that were randomly shuffled in time, space, or both. These randomized tests provide more insights into the temporal and spatial alignment of the dFC results obtained by the implemented methods.

### Spatial vs. temporal similarity of method pairs

### Comparison of actual similarities with similarity between dFC matrices with shuffled time points

### Comparison of actual similarities with similarity between dFC matrices with shuffled functional connections

### Comparison of actual similarities with similarity between dFC matrices with shuffled time points and functional connections

### Comparison of actual similarities with similarity between dFC matrices with random state time courses

## Variation over method vs. time

This section presents additional and more detailed visualizations of the analyses assessing variability of dFC results over method and time.

### Distribution of method pairs in the variation over method vs. time space

### Pair-wise variation over method vs. time scatter plots

### Group-wise variation over method vs. time scatter plots

## Variation over method vs. variation over subject

This section presents additional and more detailed visualizations of the analyses assessing variability of dFC results over method and subject.

### Variation over subject and method across functional connections

## List of Papers that have applied dFC

A selection of studies applying the seven dynamic Functional Connectivity (dFC) assessment methods included in the present study. Most studies employed only one of these methods and in most of the cases the choice was not justified further. In very few cases multiple (two or three) methods were used. The applications include clinical, and cognitive applications, as well as other applications such as the investigation of brain functional organization. These studies were found by searching the first three pages of the original papers’ cited-by list on Google Scholar and the first two pages of searching the name of the methodology on Google Scholar. Papers without available full-text versions were excluded from the search. In total, about 200 papers were reviewed, among which only 60 (43 with clinical application, 10 with cognitive application, and 7 with other applications) had actually used the examined dFC assessment methods. The rest of the papers had only cited the method’s original paper without applying it in practice. The following tables list the 60 reviewed studies along with the methods that they used and the specific application (e.g. the disease, or cognitive task).

